# SCALiR: a web application for automating absolute quantification of mass spectrometry-based metabolomics data

**DOI:** 10.1101/2023.08.16.551807

**Authors:** Stephanie L. Bishop, Luis F. Ponce-Alvarez, Soren Wacker, Ryan A. Groves, Ian A. Lewis

**Affiliations:** Department of Biological Sciences, University of Calgary, 2500 University Dr NW, Calgary, AB, Canada, T2N 1N4

## Abstract

Metabolomics is an important approach for studying complex biological systems. Quantitative liquid chromatography-mass spectrometry (LC-MS)-based metabolomics is becoming a mainstream strategy but presents several technical challenges that limit its widespread use. Computing metabolite concentrations using standard curves generated from standard mixtures of known concentrations is a labor-intensive process which is often performed manually. Currently, there are few options for open-source software tools that can automatically calculate metabolite concentrations. Herein, we introduce SCALiR (Standard Curve Application for determining Linear Ranges), a new web-based software tool specifically built for this task, which allows users to automatically transform LC-MS signal data into absolute quantitative data (https://www.lewisresearchgroup.org/software). The algorithm used in SCALiR automatically finds the equation of the line of best fit for each standard curve and uses this equation to calculate compound concentrations from their LC-MS signal. Using a standard mix containing 77 metabolites, we found excellent correlation between the concentrations calculated by SCALiR and the expected concentrations of each compound (R^2^ = 0.99) and that SCALiR reproducibly calculated concentrations of mid-range standards across ten analytical batches (average coefficient of variation 0.091). SCALiR offers users several advantages, including that it (1) is open-source and vendor agnostic; (2) requires only 10 seconds of analysis time to compute concentrations of >75 compounds; (3) facilitates automation of quantitative workflows; and (4) performs deterministic evaluation of compound quantification limits. SCALiR provides the metabolomics community with a simple and rapid tool that enables rigorous and reproducible quantitative metabolomics studies.

---

Metabolomics is a mainstream approach for studying complex biological systems, ranging from cancer^1^, infectious diseases^2^, host-microbiome interactions^3^, and microbial engineering^4,5^. The common thread among these diverse disciplines is that they all have a common need to accurately identify and quantify molecules in complex biological mixtures. Although such analyses are becoming more common in many metabolomics facilities, collecting absolute quantitative data for metabolites analyzed via routine liquid chromatography-mass spectrometry (LC-MS) analyses remains challenging^6–8^.

One of the main complications of LC-MS is that the response factor of each instrument varies day-to-day and sample-to-sample, and as a result, standard reference compounds used to calibrate signal intensities need to be acquired frequently in order to enable robust quantification^9^. These additional logistical complications of acquiring and analyzing data for standard reference materials currently make absolute quantitative metabolomics studies the exception rather than the rule and limits direct comparison between datasets obtained in different batches. Absolute quantification of metabolite signals makes it possible to directly compare data across large-cohort studies and facilitates the analysis of metabolic flux through networks^6^. Although there are many tools and resources available for metabolomics data visualization and interpretation^10^, there is a critical gap in software for performing absolute quantification of metabolomics data. Several commercial tools such as Thermo Fisher’s TraceFinder™ and open-source software packages such as Skyline for Small Molecules^11^ can determine quantitative metabolomics values from raw instrumen-tal data but require extensive user input to complete these analyses.

Most quantitative metabolomics studies conduct this work manually. This requires fitting linear curves and removing points that do not fall in the linear range, which is a time-consuming process that requires considerable analyst expertise. There are no universally accepted parameters for determining the upper and lower limits of quantification (ULOQ and LLOQ, respectively) of a compound and ensuring linearity of the curve fitting. The LLOQ is often defined by instrumental baseline noise and calculated with a numerical parameter such as 10% or a fixed multiple of the relative standard deviation of a blank matrix^12–14^. The LLOQ can also be calculated with a formula based on the percent deviation from the nominal concentration at the lowest calibrator concentration^15^. Furthermore, the process of determining the ULOQ can be complicated because numerical parameters of the curve fitting, such as homoscedasticity (equal distribution of residuals) and standard deviation of the regression, need to be considered to ensure a valid linear fit across the range of standard concentrations^16^. These challenges make the process of fitting standard curves difficult to automate and results in many investigators choosing to omit metabolite quantification from their studies.

For these reasons, there is an urgent need for software that automatically detects the upper and lower limits of quantification and the line of best fit for a series of standards and uses these data to automatically calculate the concentrations of metabolites in a sample. Herein, we introduce SCALiR (Standard Curve Application for determining Linear Ranges), a new tool designed specifically for this task. This software uses a novel algorithm that leverages a basic logarithmic property and general characteristics of LC-MS data to automatically determine the ULOQ, LLOQ, and line of best fit for external calibration standards. Using these values, SCALiR automatically transforms LC-MS peak data into absolute quantitative data. We show that SCALiR automatically generates data comparable to those obtained via manual curve fitting by an expert analyst. We then illustrate how the tool can be used to correct batch effects in large cohorts and facilitate metabolic boundary flux analysis.

## EXPERIMENTAL SECTION

### Preparation of mixed metabolite standards

All standard stock solutions were prepared from compounds ordered from Sigma-Aldrich (Oakville, ON, Canada), VWR (Edmonton, AB, Canada), or Acros Organics (now Thermo Scientific Chemicals, Waltham, MA, USA). See Table S1 for compound CAS numbers. To evaluate the linear dynamic range of each compound manually, we developed a mixed standard containing 77 compounds with variable starting concentrations and ran a 10-point dilution series with three technical replicates where each sequential standard was diluted 4-fold (1:4) with 50/50 (v/v) methanol (Fisher Optima™ LC/MS grade; Toronto, ON, Canada) and water (Fisher Optima™ LC/MS grade). Stock standard concentrations for each compound are found in the Supporting Information (Table S1).

### Liquid chromatography-mass spectrometry

All liquid chromatography-mass spectrometry (LC-MS) metabolomics data were acquired at the Calgary Metabolomics Research Facility (CMRF), according to the methods described in detail in^2,9^. Metabolite samples were resolved via a Thermo Scientific™ Vanquish™ UHPLC (Thermo Fisher Scientific) platform using hydrophilic interaction liquid chromatography (HILIC) with a 15-minute gradient. Chromatographic separation was attained using a binary solvent mixture of 20 mM ammonium formate at pH 3.0 in LC-MS grade water (Solvent A) and 0.1% formic acid (% v/v) in LC-MS grade acetonitrile (Solvent B) in conjunction with a 100 mm × 2.1 mm Syncronis™ HILIC LC column (Thermo Fisher Scientific) with a 2.1μm particle size. Data were acquired on a Thermo Scientific™ Q Exactive™ HF (Thermo Fisher Scientific) mass spectrometer in negative ionization mode.

### Manual data analysis

All LC-MS raw data files were converted to mzXML format via MSConvert GUI software^17^. LCMS analysis was conducted in El-Maven, v.0.12.0^18^ with manual peak selection using default parameters (Peak Quantitation Type: Area top; Mass Cut-off unit: 10.0 ppm). All data were then collated in Microsoft Excel, where the linear range for each compound was calculated according to visual inspection of linearity and goodness of fit. Select performance characteristics according to the U.S. FDA guidelines for bioanalytical method validation^15^ for measured compounds can be found in the Supporting Information (Table S2).

### Bacterial sample preparation and analysis

*Staphylococcus aureus* isolates were collected by the Alberta Precision Laboratories (Calgary, AB, Canada) and were prepared as described in^9^. Briefly, microbes were inoculated in Mueller-Hinton broth in 96-well plates and cultured aerobically overnight at 37°C in a humidified incubator with a 5% CO_2_ and 21% O_2_ atmosphere. After reaching an optical density between 0.35 – 0.4, culture supernatants were fixed in methanol (1:1 v/v) and centrifuged for 5 min at 4,000 x *g*. The supernatants were then diluted 1:10 with 50% methanol before LC-MS analysis.

The concentrations of metabolites in these samples were quantified using the mixed metabolite standards (see above), with each point prepared as an 8-point dilution series. The highest concentration standard was prepared as a 2-fold (1:2) dilution in 50/50 (v/v) methanol (Fisher Optima™ LC/MS grade) and water (Fisher Optima™ LC/MS grade) from the stock standard concentrations described in Table S1. The next seven dilutions were prepared as 4-fold (1:4) dilutions in the same 50% methanol solvent. LC-MS data were acquired as described above. Two 100 mm × 2.1 mm Syncronis™ HILIC LC columns (Thermo Fisher Scientific) were used to analyze multiple batches. All data were quantified manually as described above and compared to automated data quantification via SCALiR.

### Implementation, statistical analysis, and visualization

The SCALiR backend was implemented in Python (Version 3) using the standard libraries pandas, numpy, and matplotlib. The user interface was created on Streamlit and the source code is available in GitHub (https://github.com/LewisResearchGroup/ms-conc). All statistical analysis of data was performed in Python (Version 3) or GraphPad Prism (Version 9.5.1). Figures were created using Python, GraphPad Prism, Adobe Illustrator (Version 27.2), Inkscape (Version 1.2), and by directly downloading images and screenshots from the SCALiR app.

### Safety considerations

Reagents used in this investigation do not pose any significant safety risks outside of those experienced regularly when working with moderate strength acids and solvents.

## RESULTS AND DISCUSSION

### SCALiR algorithm

One of the principal challenges in quantitative metabolomics is the lack of a standardized or deterministic method for computing the LLOQ and ULOQ for a series of standards. The algorithm incorporated into SCALiR takes advantage of a basic logarithmic property and an intrinsic characteristic of LC-MS signals that allows the curve fitting process to be generalized. Specifically, in linear datasets where the slope is much greater than the intercept, the log-log transform of the linear series will have a slope approaching one. This transformation is detailed in Eq. 1-6 below.

Robust calibration reference standards follow a relationship wherein signal intensities (*y*) and metabolite concentrations (*x*) are described by a linear model:

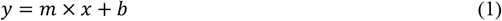

where *m* denotes slope and *b* denotes the y-intercept. This relationship can be transformed as follows:

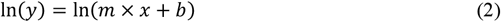

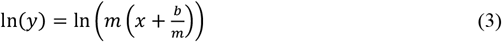

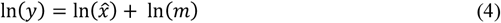

where 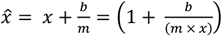

In cases where *b* ≪(*m* ×*x*), in Eq. (4), 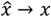 and becomes:

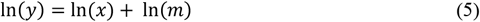

which can be re-written as:

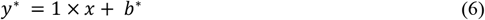

where *y*^*^ is the LC-MS signal in the *ln* scale, the slope is equal to 1, and *b*^*^ is ln(*m*).

To illustrate this point, we calculated the intercept fraction value, 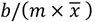, for ten metabolites across ten batches using the average concentration of the five lowest concentration standards of an 8-point standard curve to ensure points fell below the ULOQ (excluding points with no signal). As shown in Figure 1a, these values fall within the range (-0.1, 0.1) and this distribution range holds for 80% of the metabolites we measured (Figure 1b). When *b* ≪(*m* ×*x*), the curve in the linear scale (Eq. 1) is transformed into a linear curve in the *ln* scale with a slope equal to 1 and the intercept equal to the *ln* of the slope in the linear scale (Figure 1c,d). Compounds that exhibit deviations from this linear relationship, which result in a less adequate fit after the *ln* transformation with a slope not close to 1 and an intercept fraction value ≫ 0 (examples shown in Figure S1), are automatically flagged as a “failed fit” by SCALiR.

**Figure 1.**
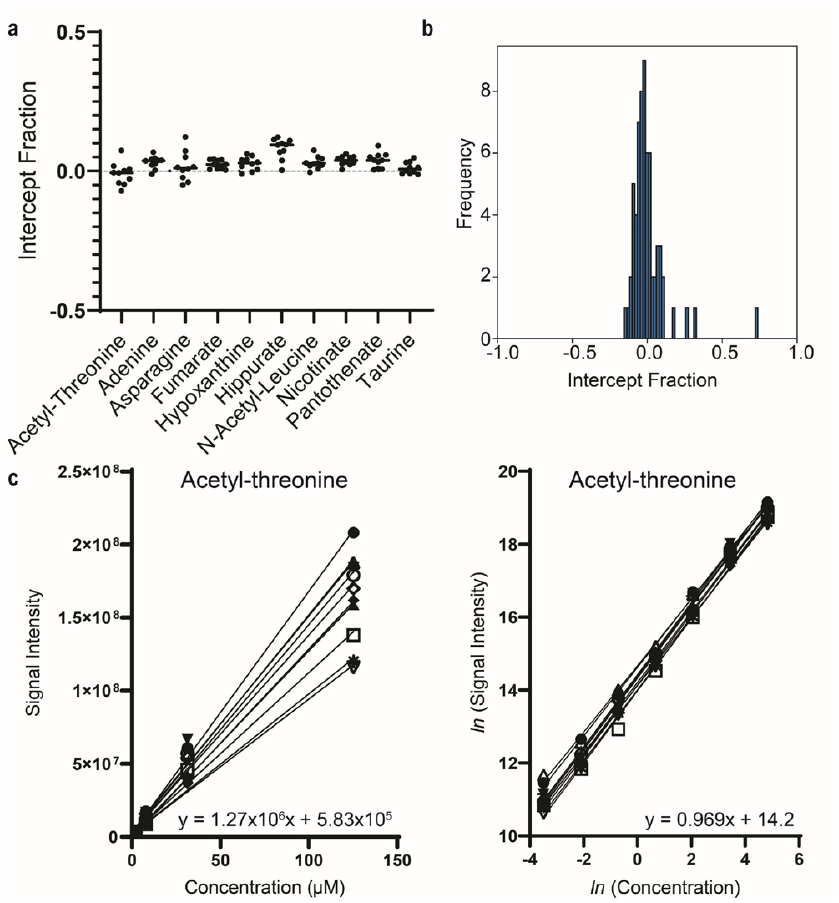
(a) Examples of intercept fraction values (*b*/(*m* ×*x* )) for the standard curves of ten metabolites run over ten batches. (b) The distribution of intercept fraction values for standard curves of all metabolites measured. (c) Examples of a set of ten standard curves for acetyl-threonine in the linear and *ln* scale. Each curve corresponds to one batch and the line of best fit equations show an example of the slope and y-intercept from one standard curve.

The algorithm used by SCALiR incorporates these properties to determine the ULOQ and LLOQ for the standard curve, as well as an iterative fitting process to find the line of best fit. The goodness of fit for the standard curve is controlled by the residual function (*r*) between the expected (*e*) and observed (*o*) values defined as:

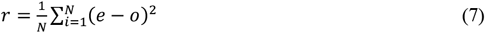

where *N* is the number of points in the dataset.

When *r* is calculated to be above the experimentally determined threshold value of 0.01 (Figure S2), the point with the largest distance between the predicted and the experimental value is removed from the dataset (Figure 2). This step is repeated until the value of *r* falls below the set threshold, indicating acceptable goodness of fit.

**Figure 2.**
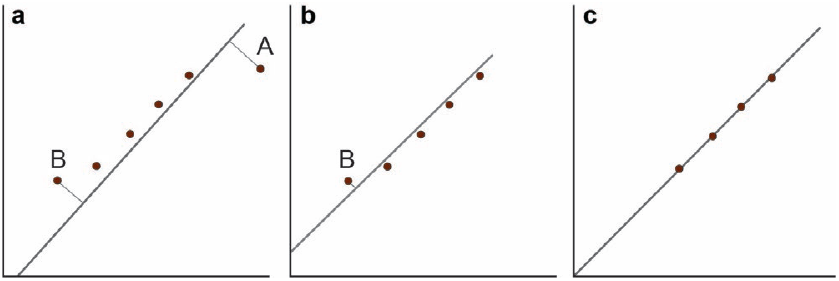
Iterative regression algorithm for finding the linear range of a series of standards. (a) The best fit curve is generated from the dataset. (b) When the residual (*r*) of the curve is > 0.01, the farthest point from the curve (“A”) is removed and repeated (“B”). This step is carried out again until (c) the data allow a linear regression fitting that achieves the quality threshold for *r*.

### SCALiR app design

The web-based application SCALiR (https://www.lewisresearchgroup.org/software) is designed to work with peak data files generated from open-source software El-Maven^18^ or MINT (https://mint.resistancedb.org/) without further modifications. Peak data from any software can be used if the peak data file follows the formatting in the sample files accessed directly from the app. Sample data upload files for standards concentrations can also be accessed directly from the app. Instructions on how to use the app are explained on the web application interface and a tutorial with demo data files for upload can be accessed via the web interface and are included in the Supporting Information. Users can also document issues or suggestions for the app at https://github.com/LewisResearchGroup/ms-conc/issues.

Figure 3 shows key steps and features of the app. Users upload a comma-separated values (CSV) file containing concentrations of each standard as well as a CSV or Excel file with peak list information for standards and samples (Figure 3a). The user then selects which program was used to generate the peak list information and chooses the fixed slope, wide slope, or interval slope option, and can adjust the permissible values of the slope for the interval slope option (Figure 3b). Once the program is run, the user can download the standard curve parameters, including slope, intercept, and linear range minimum and maximum for each compound, as well as the calculated concentrations for standards and samples. Additionally, the user can view log-log plots showing the standard curve for each compound and download individual images of each plot with the option to include axis labels or leave them blank before downloading the images.

**Figure 3.**
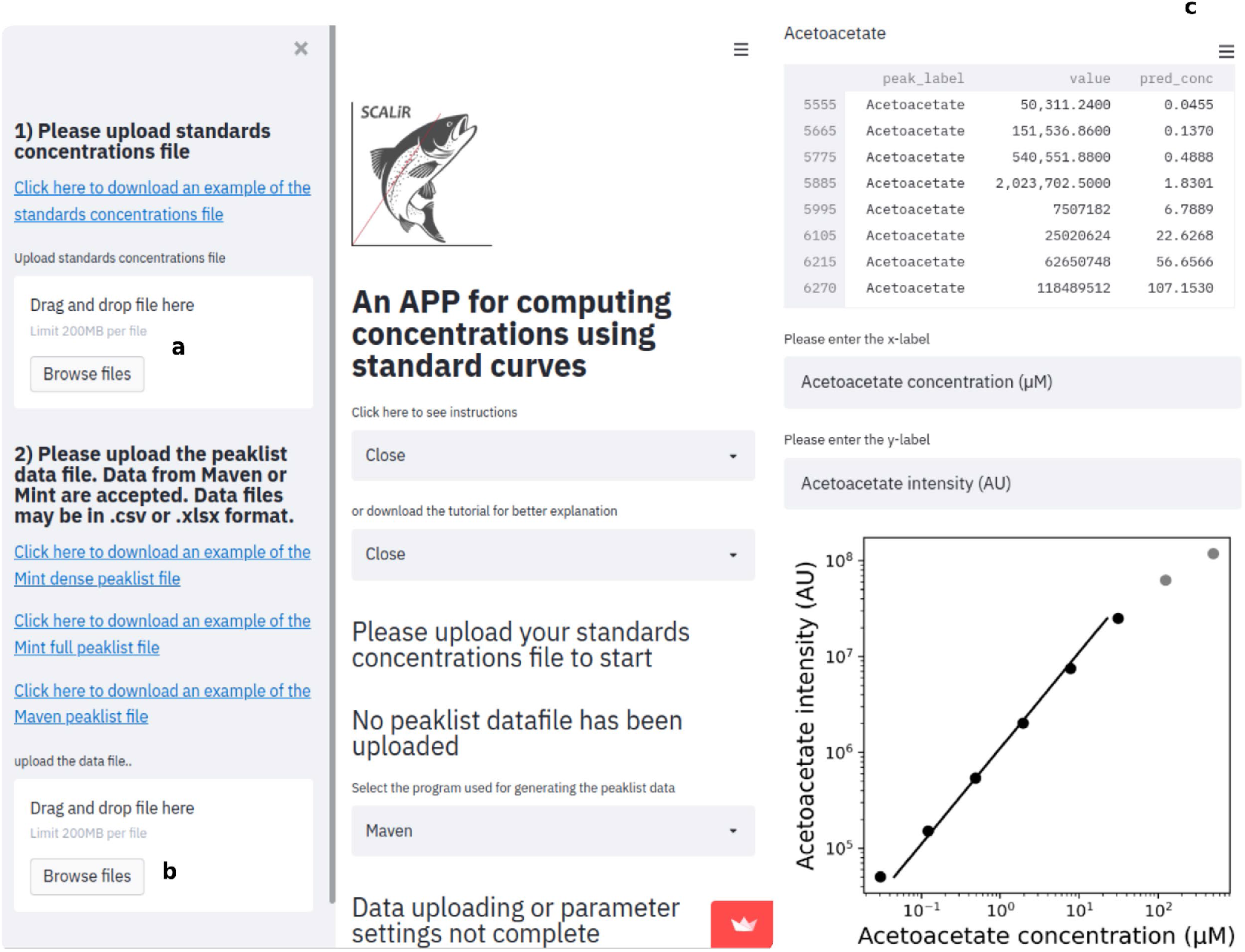
Features of the web-based application SCALiR and schematic of its iterative standard curve fitting process. Users upload standard concentrations (a) and peak list data files (b) and select settings for generating the standard curves. (c) SCALiR performs its iterative fitting algorithm, stopping when the fitting reaches the residual threshold (*r* ≤ 0.01). The remaining maximum and minimum concentration values are reported as the range for the linear behavior. Users can download the results for standard curve parameters, concentration data, and visualize individual standard curves as log-log plots.

### Performance evaluation

To assess the performance of SCALiR, we compared the concentrations calculated by the app to the actual concentration values of each metabolite in an 8-point standard dilution series (Figure 4a; supplementary datafile). The data shown here are from the mixed standard containing 77 metabolites and all validation data are shown with SCALiR’s fixed slope feature. Similar results were obtained when we used the wide slope option. The concentration values calculated by the app correlated closely with the actual concentration values of the standards (R^2^ = 0.99 in the *ln* scale; Figure 4a), demonstrating that SCALiR’s algorithm accurately determines the linear range and concentrations of metabolites. This value was comparable to the correlation between calculated and actual concentrations obtained by an expert analyst that manually fitted standard curves (R^2^ = 0.98 in the *ln* scale; Figure 4b). Thirdly, we compared SCALiR against concentration values generated by an expert analyst manually (Figure 4c). Again, we found excellent correlation between the concentration values calculated by SCALiR and those calculated by manual inspection (R^2^ = 0.99 in the *ln* scale; Figure 4c), indicating that SCALiR’s algorithm performs very similarly to a trained analyst.

**Figure 4.**
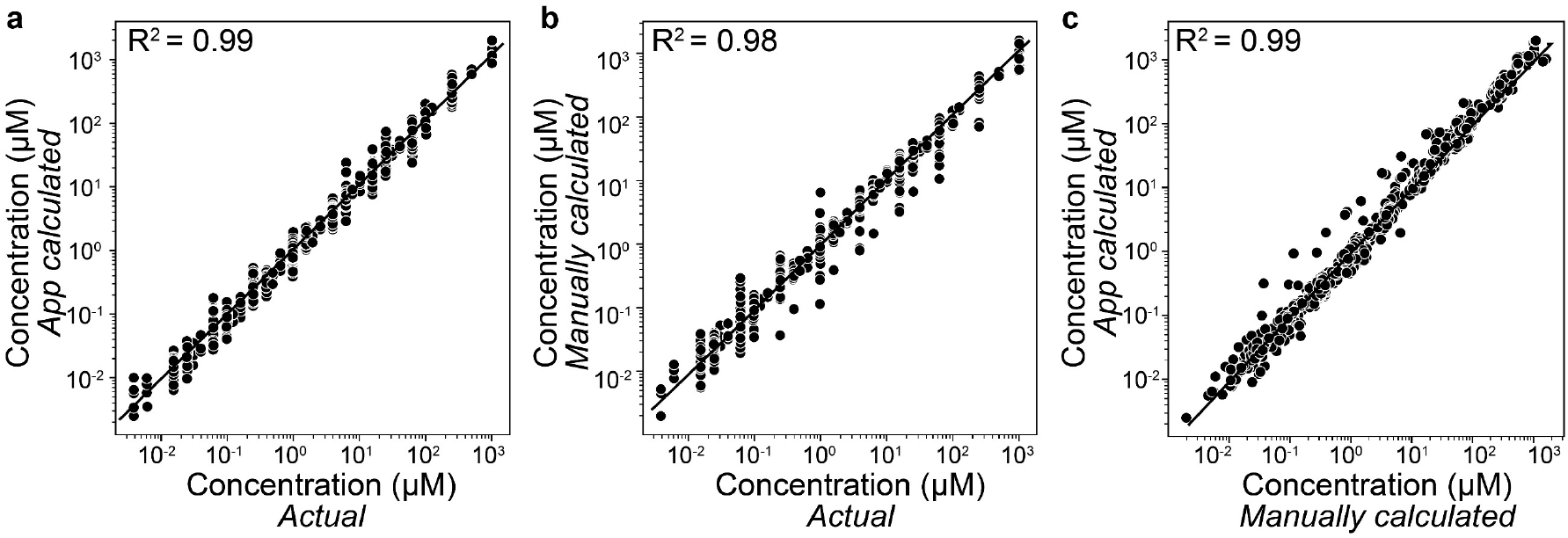
Validation of the algorithm used in SCALiR for calculating concentrations of compounds in a standard mix. (a) Comparison between the app calculated concentrations in standards and the actual concentrations for each compound. (b) Comparison between the manually calculated concentrations and actual concentrations for each compound. (c) Comparison between the app calculated concentrations in the standard samples and manually calculated concentrations.

SCALiR also demonstrated reproducible results across ten separate analytical batches (Table S3). We calculated that the average coefficient of variation (CV) for the middle four standards in the 8-point standard curves (Standards 3 – 6) of ten representative compounds was between 0.054 – 0.152, with an average CV of 0.091 for all ten compounds. Additionally, 85% of the SCALiR-calculated concentrations of these standards met the US FDA guidelines stating that non-zero calibrators should be within ±15% of the nominal (expected) concentration^15^ (Table S3).

### Analytical challenges solved by SCALiR

One of the major advantages of using SCALiR is its deterministic process for evaluating the linear range of a standard series, which is based on empirically determined parameters. Manual evaluation of quantification limits is not only time-consuming but can lead to variation in data analysis results between analysts and over separate batches. This can result in differences in quantification limits and affect the accuracy of the sample concentrations calculated. Figure 5 shows an example of challenges that can occur when a standard series displays multiple linear regimes, or regions of the curve that display a linear fit with a coefficient of determination (R^2^) > 0.99. Using hippurate as an example, we show that an 8-point standard series transformed to the *ln* scale can display two or more regions that would have a valid linear fit based on the coefficient of determination parameter (Figure 5a). However, only two of these regions satisfy the linear relationship between signal intensity and concentration required for the valid *ln* transformation showing a slope of the regression curve equal to approximately 1 (Lower curve in Figure 5a).

**Figure 5.**
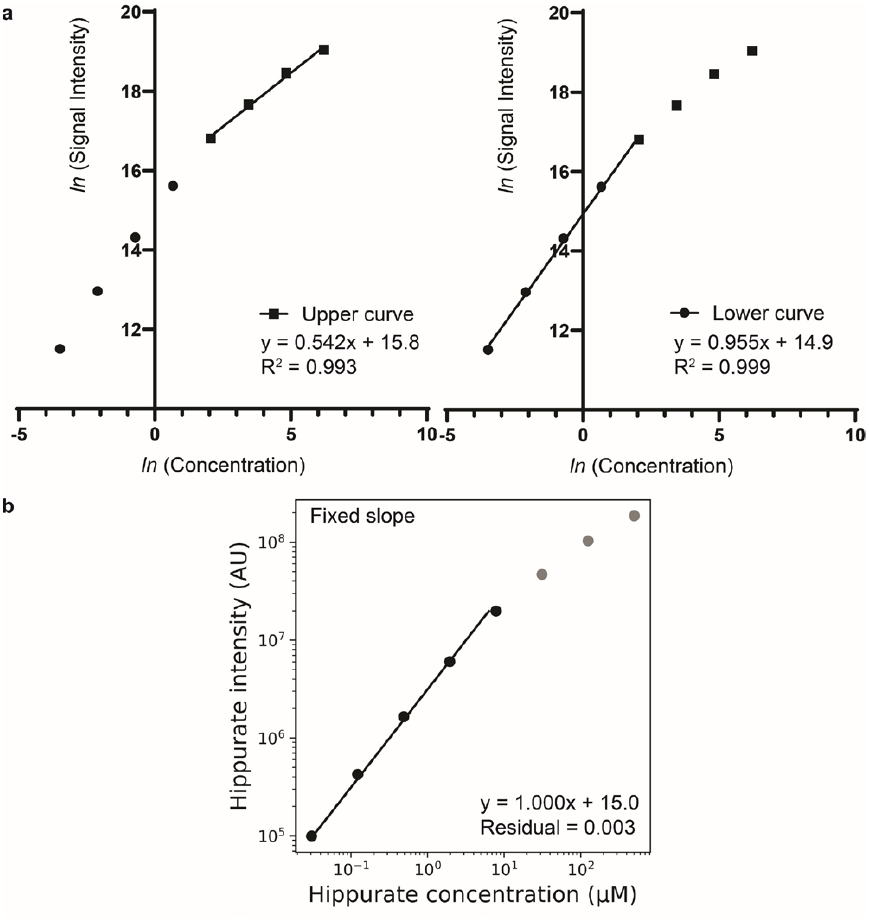
SCALiR simplifies the process for determining the linear range of a standard series where there are two or more distinct linear regimes. The 8-point standard curve for hippurate in the natural logarithmic scale is shown as an example in (a), which can result in two possible linear regions with a high R^2^ value (>0.99). (b) Log-log plot downloaded from SCALiR, which uses a deterministic fitting algorithm that sets the slope of the regression curve to 1.000 in the *ln* scale with the fixed slope option and stops the fitting when the residual of the curve < 0.01.

SCALiR automatically determines the region of the curve that both satisfies the linear relationship requirement (slope of the *ln* transformed curve = 1) and provides a high-quality fit (residual of the curve < 0.01) (Figure 5b). An example of a log-log plot downloaded from the app shows the standard curve of an individual metabolite, hippurate. In summary, SCALiR’s deterministic algorithm standardizes the process of determining a linear range for a standard series when multiple valid linear fittings are possible, and thus can lead to more reproducible quantitative results in a fraction of the analysis time of traditional methods.

### Applications

SCALiR can alleviate common quantitative LC-MS challenges including batch effects and has clear utility for a variety of quantitative applications, including analyzing cellular boundary flux^19–21^ - an approach which is gaining traction as a foundation for clinical diagnostics^2^, analyzing metabolic interactions within microbial communities^22,23^, and assessing the microbial production of biofuels^24,25^. Batch effects are mainly caused by instrumental drift in signal over time and exacerbated by using varied analytical conditions (e.g., different chromatography columns or solvent batches)^9^. In this example, we show how SCALiR can minimize batch effects arising from a multiplexed strategy that uses two separate chromatography columns to enable high-throughput analysis of over 3,000 injections of the same sample of *Staphylococcus aureus* growth medium^9^ (Figure S3). Clear batch effects were apparent when we compared only the signal intensity values for arginine in a dataset (Figure 6a). Using SCALiR to calculate concentrations of arginine in each batch, we generated a stable result across the repeated injections of the sample irrespective of the column used (Figure 6b).

**Figure 6.**
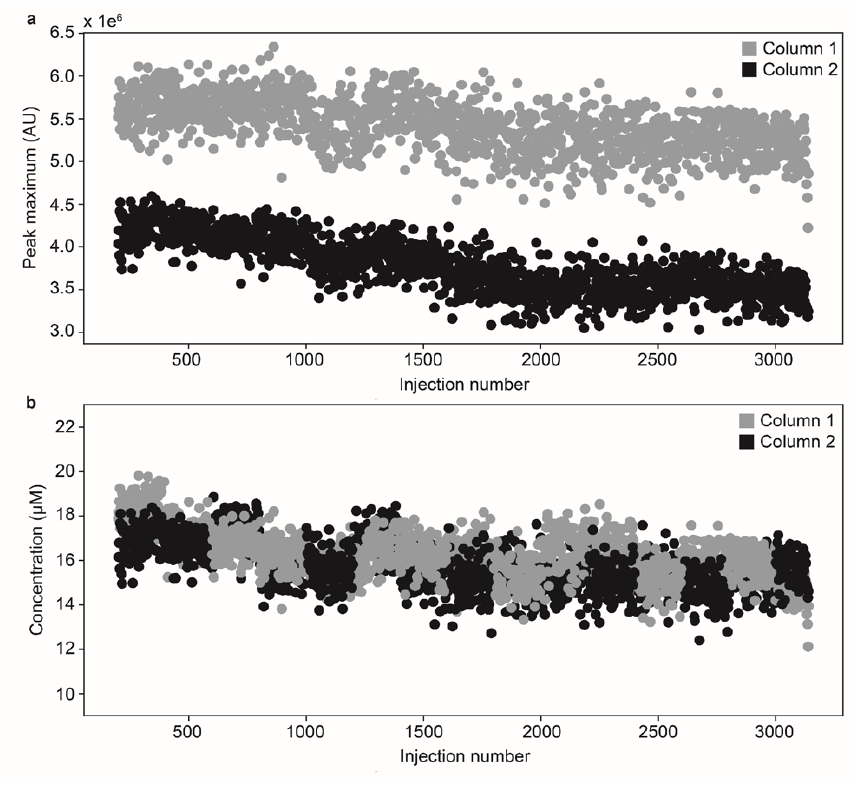
Comparison of arginine (a) peak maximum values (arbitrary units) and (b) concentration values (μM) in samples across two columns in a large-scale Staphylococcus aureus isolates dataset with a multiplexed chromatography strategy.

Additionally, we used SCALiR to perform a metabolic boundary flux analysis of the Mueller Hinton growth medium components consumed by *S. aureus* in the large-cohort study described previously (Figure 7). Using our quantitative approach, we revealed different metabolic phenotypes that were not discernible using LC-MS signal data alone. For example, we found that *S. aureus* cells mostly consume serine, arginine, and trehalose (Figure 7b) as opposed to using other carbon and nitrogen sources calculated from the normalized peak area (Figure 7a). Using the quantitative approach, we also observed that some metabolites such as glutamate and aspartate, despite their high availability, are not preferentially consumed by *S. aureus* (Figure 7c), which suggests a route to growth medium optimization. This approach also allowed us to infer the main metabolic routes used by the cells for optimal growth (Figure 7d). By facilitating quantitative metabolomics studies, SCALiR represents an important new tool that may soon serve as a new standard for a wide range of metabolomics applications, including biomarker discovery for a variety of diseases such as Alzheimer’s disease^26^ and cancer^27^, as well as studies of plant secondary metabolism^28^.

**Figure 7.**
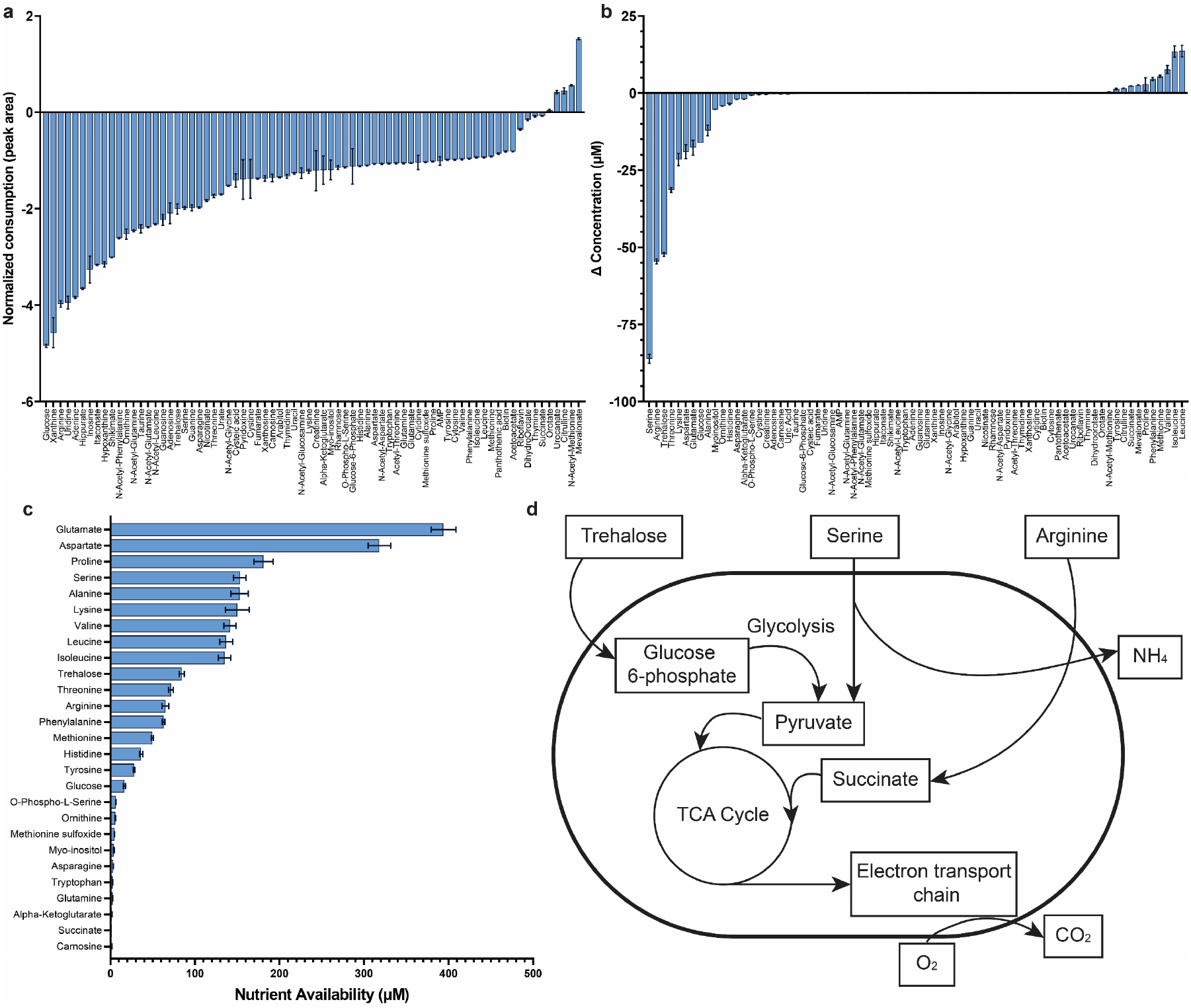
Boundary flux analysis of *S. aureus* clinical isolates grown in Mueller Hinton (MH) growth medium. (a) Metabolite consumption/secretion assessment by the ratio of peak intensities in the microbial supernatant and MH medium. (b) Metabolite consumption/secretion analysis using concentration differences. (c) Nutrient availability in MH medium. (d) Proposed metabolic pathways for optimal *S. aureus* growth, according to nutrient consumption and secretion data.

### Limitations

One limitation of SCALiR is that it can only compute the line of best fit for linear data and uses a logarithmic transformation which may not be appropriate for all datasets^16^. Users may, however, choose to fit their data with a non-linear fitting using the interval slope or wide slope options available in the app. Additionally, as with any method used to calculate concentrations from a standard curve, SCALiR can only provide accurate values for concentrations in the respective linear range for a given compound. With these considerations in mind, SCALiR provides users with a new tool to generate quantitative metabolomics data over a wide range of distinct compounds and concentration ranges.

## CONCLUSION

Quantitative metabolomics is rapidly emerging as an important component of many metabolomics research programs; however, the complexity and time required to quantify metabolites has limited its widespread use. SCALiR provides users with three main advantages compared to the traditional method of manually fitting standard curves for quantitative metabolomics: (1) reduced analysis time, requiring only 10 seconds to compute concentrations of >75 compounds; (2) automation of quantitative workflows with minimal training or computational expertise required; and (3) deterministic evaluation of compound quantification limits and line of best fit, reducing analytical subjectivity. Additionally, SCALiR is open-source and vendor agnostic. SCALiR fits an important need in the metabolomics community and opens the door to routine quantitative metabolomics workflows.

## Supporting information

Tutorial

Sample Data 1

Sample Data 2

Sample Data 3

Sample Data 4

## ASSOCIATED CONTENT

### Supporting Information

The Supporting Information is available free of charge on the

ACS Publications website.

Supporting figures and tables (PDF)

Bacterial metabolomics concentrations comparison manual vs

SCALiR data file (CSV)

Bacterial metabolomics samples and standards SCALiR concentrations data file (CSV)

Raw signal intensity data manual standard curves Maven format peak list negative ionization mode (CSV)

Raw signal intensity data manual standard curves Maven format peak list positive ionization mode (CSV)

SCALiR Tutorial (PDF)

SCALiR Standards Concentrations File (CSV)

SCALiR MINT Peaklist Dense Peak Max (CSV)

SCALiR MINT Peaklist Full Results (CSV)

SCALiR Maven Peaklist (CSV)

## Data availability

All data are available in the Supporting Information of this manuscript or at https://pubs.acs.org/doi/10.1021/acs.analchem.2c00078?ref=PDF. SCALiR source code is available at https://github.com/LewisResearchGroup/ms-conc.

## AUTHOR INFORMATION

### Corresponding Author

^*^Ian A. Lewis

### Author Contributions

IAL and LFPA conceived of the project. LFPA and SW developed the algorithm and web application. SLB and RAG performed all experimental analyses. LFPA and SLB performed all data analysis and SLB wrote the primary draft of the manuscript. All authors have given approval to the final version of the manuscript.

^‡^These authors contributed equally.

## ACKNOWLEDGMENT

We thank K. A. Pittman and N. van Bavel for their critical review and editorial help in preparing this article. IAL is supported by an Alberta Innovates — Health Solutions (AIHS, Translational Health Chair), a 2016 Genome Canada Genomics Applied Partnership Program (GAPP) and a 2017 Genome Canada Large Scale Applied Research Project award (both administered by Genome Alberta), the Natural Sciences and Engineering Research Council (NSERC) [DG 04547], the Canada Foundation for Innovation (CFI; JELF-34986), and the International Microbiome Centre and IMPACTT Microbiome Research Core (CIHR IMC-161484). SLB is supported by a National Institutes of Health (NIH) grant [Project Number 1R01AI153521-01] and a Canadian Institutes of Health Research (CIHR) Postdoctoral Fellowship [Project Number 202110HIV-477488-87373]. SW is directly supported via a Genome Canada 2017 Large Scale Applied Research Project award administered through Genome Alberta.

## REFERENCES

(1) Schmidt, D. R.; Patel, R.; Kirsch, D. G.; Lewis, C. A.; Vander Heiden, M. G.; Locasale, J. W. Metabolomics in Cancer Research and Emerging Applications in Clinical Oncology. CA. Cancer J. Clin. 2021, 71 (4), 333–358. https://doi.org/10.3322/caac.21670.

(2) Rydzak, T.; Groves, R. A.; Zhang, R.; Aburashed, R.; Pushpker, R.; Mapar, M.; Lewis, I. A. Metabolic Preference Assay for Rapid Diagnosis of Bloodstream Infections. Nat. Commun. 2022, 13 (1), 1–12. https://doi.org/10.1038/s41467-022-30048-6.

(3) Bishop, S. L.; Drikic, M.; Wacker, S.; Chen, Y. Y.; Kozyrskyj, L.; Lewis, I. A. Moving beyond Descriptive Studies: Harnessing Metabolomics to Elucidate the Molecular Mechanisms Underpinning Host-Microbiome Phenotypes. Mucosal Immunol. 2022, 15 (6), 1071–1084. https://doi.org/10.1038/s41385-022-00553-4.

(4) Jang, Y. S.; Woo, H. M.; Im, J. A.; Kim, I. H.; Lee, S. Y. Metabolic Engineering of Clostridium Acetobutylicum for Enhanced Production of Butyric Acid. Appl. Microbiol. Biotechnol. 2013, 97 (21), 9355–9363. https://doi.org/10.1007/s00253-013-5161-x.

(5) Martien, J. I.; Amador-Noguez, D. Recent Applications of Metabolomics to Advance Microbial Biofuel Production. Curr. Opin. Biotechnol. 2017, 43, 118–126. https://doi.org/10.1016/j.copbio.2016.11.006.

(6) Alseekh, S.; Aharoni, A.; Brotman, Y.; Contrepois, K.; D’Auria, J.; Ewald, J.; C. Ewald, J.; Fraser, P. D.; Giavalisco, P.; Hall, R. D.; Heinemann, M.; Link, H.; Luo, J.; Neumann, S.; Nielsen, J.; Perez de Souza, L.; Saito, K.; Sauer, U.; Schroeder, F. C.; Schuster, S.; Siuzdak, G.; Skirycz, A.; Sumner, L. W.; Snyder, M. P.; Tang, H.; Tohge, T.; Wang, Y.; Wen, W.; Wu, S.; Xu, G.; Zamboni, N.; Fernie, A. R. Mass Spectrometry-Based Metabolomics: A Guide for Annotation, Quantification and Best Reporting Practices. Nat. Methods 2021, 18 (7), 747–756. https://doi.org/10.1038/s41592-021-01197-1.

(7) Lu, W.; Su, X.; Klein, M. S.; Lewis, I. A.; Fiehn, O.; Rabinowitz, J. D. Metabolite Measurement: Pitfalls to Avoid and Practices to Follow. Annu. Rev. Biochem. 2017, 86, 277– 304. https://doi.org/10.1146/annurev-biochem-061516-044952.

(8) Kapoore, R. V.; Vaidyanathan, S. Towards Quantitative Mass Spectrometry-Based Metabolomics in Microbial and Mammalian Systems. Philos. Trans. R. Soc. A Math. Phys. Eng. Sci. 2016, 374 (2079). https://doi.org/10.1098/rsta.2015.0363.

(9) Groves, R. A.; Mapar, M.; Aburashed, R.; Ponce, L. F.; Bishop, S. .; Rydzak, T.; Drikic, M.; Bihan, D. G.; Benediktsson, H.; Clement, F.; Gregson, D. B.; Lewis, I. A. Methods for Quantifying the Metabolic Boundary Fluxes of Cell Cultures in Large Cohorts by High-Resolution Hydrophilic Liquid Chromatography Mass Spectrometry. Anal. Chem. 2022, 1–9.

(10) Misra, B. B. New Software Tools, Databases, and Resources in Metabolomics: Updates from 2020. Metabolomics 2021, 17 (5), 1–24. https://doi.org/10.1007/s11306-021-01796-1.

(11) Adams, K. J.; Pratt, B.; Bose, N.; Dubois, L. G.; St. John-Williams, L.; Perrott, K. M.; Ky, K.; Kapahi, P.; Sharma, V.; Maccoss, M. J.; Moseley, M. A.; Colton, C. A.; Maclean, B. X.; Schilling, B.; Thompson, J. W. Skyline for Small Molecules: A Unifying Software Package for Quantitative Metabolomics. J. Proteome Res. 2020, 19 (4), 1447–1458. https://doi.org/10.1021/acs.jproteome.9b00640.

(12) Thomson, M.; Ellison, S. L. R.; Wood, R. Harmonized Guidelines for Single-Laboratory Validation of Methods of Analysis (IUPAC Technical Report). Pure Appl. Chem. 2002, 74 (5), 835–855.

(13) EPA. Definition and Procedure for the Determination of the Method Detection Limit, Revision 2; 2016.

(14) Betz, J. M.; Brown, P. N.; Roman, M. C. Accuracy, Precision, and Reliability of Chemical Measurements in Natural Products Research. Fitoterapia 2011, 82 (1), 44–52. https://doi.org/10.1016/j.fitote.2010.09.011.

(15) U.S. FDA. Guidance for Industry Bioanalytical Method Validation Guidance for Industry Bioanalytical Method Validation; 2018.

(16) Jurado, J. M.; Alcázar, A.; Muñiz-Valencia, R.; Ceballos-Magaña, S. G.; Raposo, F. Some Practical Considerations for Linearity Assessment of Calibration Curves as Function of Concentration Levels According to the Fitness-for-Purpose Approach. Talanta 2017, 172 (May), 221–229. https://doi.org/10.1016/j.talanta.2017.05.049.

(17) Chambers, M. C.; MacLean, B.; Burke, R.; Amodei, D.; Ruderman, D. L.; Neumann, S.; Gatto, L.; Fischer, B.; Pratt, B.; Egertson, J.; Hoff, K.; Kessner, D.; Tasman, N.; Shulman, N.; Frewen, B.; Baker, T. A.; Brusniak, M. Y.; Paulse, C.; Creasy, D.; Flashner, L.; Kani, K.; Moulding, C.; Seymour, S. L.; Nuwaysir, L. M.; Lefebvre, B.; Kuhlmann, F.; Roark, J.; Rainer, P.; Detlev, S.; Hemenway, T.; Huhmer, A.; Langridge, J.; Connolly, B.; Chadick, T.; Holly, K.; Eckels, J.; Deutsch, E. W.; Moritz, R. L.; Katz, J. E.; Agus, D. B.; MacCoss, M.; Tabb, D. L.; Mallick, P. A Cross-Platform Toolkit for Mass Spectrometry and Proteomics. Nat. Biotechnol. 2012, 30 (10), 918–920. https://doi.org/10.1038/nbt.2377.

(18) Melamud, E.; Vastag, L.; Rabinowitz, J. D. Metabolomic Analysis and Visualization Engine for LC - MS Data. Anal. Chem. 2010, 82 (23), 9818–9826. https://doi.org/10.1021/ac1021166.

(19) Hui, S.; Cowan, A. J.; Zeng, X.; Yang, L.; TeSlaa, T.; Li, X.; Bartman, C.; Zhang, Z.; Jang, C.; Wang, L.; Lu, W.; Rojas, J.; Baur, J.; Rabinowitz, J. D. Quantitative Fluxomics of Circulating Metabolites. Cell Metab. 2020, 32 (4), 676-688.e4. https://doi.org/10.1016/j.cmet.2020.07.013.

(20) Orth, J. D.; Thiele, I.; Palsson, B. O. What Is Flux Balance Analysis? Nat. Biotechnol. 2010, 28 (3), 245–248. https://doi.org/10.1038/nbt.1614.

(21) Pinu, F. R.; Villas-Boas, S. G.; Aggio, R. Analysis of Intracellular Metabolites from Microorganisms: Quenching and Extraction Protocols. Metabolites 2017, 7 (4). https://doi.org/10.3390/metabo7040053.

(22) Khandelwal, R. A.; Olivier, B. G.; Röling, W. F. M.; Teusink, B.; Bruggeman, F. J. Community Flux Balance Analysis for Microbial Consortia at Balanced Growth. PLoS One 2013, 8 (5). https://doi.org/10.1371/journal.pone.0064567.

(23) Perez-Garcia, O.; Lear, G.; Singhal, N. Metabolic Network Modeling of Microbial Interactions in Natural and Engineered Environmental Systems. Front. Microbiol. 2016, 7 (MAY). https://doi.org/10.3389/fmicb.2016.00673.

(24) Hollinshead, W.; He, L.; Tang, Y. J. Biofuel Production: An Odyssey from Metabolic Engineering to Fermentation Scale-Up. Front. Microbiol. 2014, 5 (JULY), 1–8. https://doi.org/10.3389/fmicb.2014.00344.

(25) Carnicer, M.; Vieira, G.; Brautaset, T.; Portais, J. C.; Heux, S. Quantitative Metabolomics of the Thermophilic Methylotroph Bacillus Methanolicus. Microb. Cell Fact. 2016, 15 (1), 1–12. https://doi.org/10.1186/s12934-016-0483-x.

(26) Varma, V. R.; Oommen, A. M.; Varma, S.; Casanova, R.; An, Y.; Andrews, R. M.; O’Brien, R.; Pletnikova, O.; Troncoso, J. C.; Toledo, J.; Baillie, R.; Arnold, M.; Kastenmueller, G.; Nho, K.; Doraiswamy, P. M.; Saykin, A. J.; Kaddurah-Daouk, R.; Legido-Quigley, C.; Thambisetty, M. Brain and Blood Metabolite Signatures of Pathology and Progression in Alzheimer Disease: A Targeted Metabolomics Study; 2018; Vol. 15. https://doi.org/10.1371/journal.pmed.1002482.

(27) Sullivan, M. R.; Danai, L. V.; Lewis, C. A.; Chan, S. H.; Gui, D. Y.; Kunchok, T.; Dennstedt, E. A.; Heiden, M. G. V.; Muir, A. Quantification of Microenvironmental Metabolites in Murine Cancers Reveals Determinants of Tumor Nutrient Availability. Elife 2019, 8, 1–27. https://doi.org/10.7554/eLife.44235.

(28) Jorge, T. F.; Mata, A. T.; António, C. Mass Spectrometry as a Quantitative Tool in Plant Metabolomics. Philos. Trans. R. Soc. A Math. Phys. Eng. Sci. 2016, 374 (2079). https://doi.org/10.1098/rsta.2015.0370.

