## Supporting Information for "SCALiR: a web application for automating absolute quantification of mass spectrometry-based metabolomics data"

Table S1. Compound information and highest standard concentration for mixed metabolite standard dilution series.

| Chemical (pure compound) | Chemical Name | Manufacturer/ CAS No. | Average m/z | Average Retention Time (min) | Stock Concentration in Mixed Standard (µM) |
|---|---|---|---|---|---|
| (R)-Mevalonate | (R)-Mevalonic acid lithium salt | 1255502-07-8 (Sigma-Aldrich) | 147.06626 | 2.0093 | 101.24 |
| 2-Oxoglutarate | Alpha-ketoglutaric acid | 305-72-6 (Sigma-Aldrich) | 145.01422 | 8.4733 | 999.68 |
| Acetoacetate | Lithium acetoacetate | 3483-11-2 (Sigma-Aldrich) | 101.02433 | 0.8953 | 997.96 |
| Acetyl-Threonine | Acetyl-L-Threonine | 17093-74-2 (Santa Cruz Biotechnology) | 160.06154 | 4.3750 | 1002.64 |
| Adenine | Adenine | 73-24-5 (Sigma-Aldrich) | 134.04717 | 4.8400 | 101.38 |
| Adenosine | Adenosine | 58-61-7 (Sigma-Aldrich) | 266.08950 | 4.4190 | 99.91 |
| Adenosine 5'-monophosphate | Adenosine 5-monophosphate monohydrate | 18422-05-4 (Sigma-Aldrich) | 346.05585 | 8.5283 | 99.11 |
| Biotin | D-biotin | 58-85-5 (VWR) | 243.08081 | 3.6207 | 100.69 |
| Carnosine | L-Carnosine | 305-84-0 (Sigma-Aldrich) | 225.09925 | 8.6820 | 99.90 |
| Creatinine | Creatinine | 60-27-5 (Sigma-Aldrich) | 112.05163 | 5.5830 | 101.66 |
| Cytidine | Cytidine | 65-46-3 (Sigma-Aldrich) | 242.0782 | 6.143 | 98.68 |
| D-Arabitol | D-(+)-Arabitol | 488-82-4 (Sigma-Aldrich) | 151.06123 | 4.6943 | 996.39 |
| D-Fructose 1-6 Bisphosphate | D-fructose 1,6-bisphosphate trisodium salt hydrate | 38099-82-0 (Sigma-Aldrich) | 338.98884 | 9.9490 | 999.85 |
| D-Glucose | D-(+)-Glucose | 50-99-7 (Sigma-Aldrich) | 179.05605 | 5.2140 | 998.00 |

| Chemical (pure compound) | Chemical Name | Manufacturer/ CAS No. | Average m/z | Average Retention Time (min) | Stock Concentration in Mixed Standard (µM) |
|---|---|---|---|---|---|
| D-Glucose-6-Phosphate | D-glucose 6-phosphate sodium salt | 54010-71-8 (Sigma-Aldrich) | 259.02234 | 8.7567 | 998.16 |
| DL-Homocysteine | DL-homocysteine | 454-29-5 (Sigma-Aldrich) | 134.02810 | 7.3323 | 99.72 |
| Folate | Folate | 59-30-3 (Sigma-Aldrich) | 440.13249 | 7.7190 | 40.60 |
| Fumarate | Sodium fumarate dibasic | 17013-01-3 (Sigma-Aldrich) | 115.00365 | 4.0853 | 1001.00 |
| Glycine | Glycine | 56-40-6 (Sigma-Aldrich) | 74.02479 | 8.0510 | 1004.40 |
| Guanine | Guanine | 73-40-5 (Sigma-Aldrich) | 150.04216 | 5.9087 | 99.91 |
| Guanosine | Guanosine | 118-00-3 (Sigma-Aldrich) | 282.08441 | 6.2460 | 100.27 |
| Hippurate | Sodium hippurate hydrate | 532-94-5 (Sigma-Aldrich) | 178.05084 | 1.9217 | 1003.23 |
| Hypoxanthine | Hypoxanthine | 68-94-0 (Sigma-Aldrich) | 135.03120 | 4.5713 | 97.72 |
| Inosine | Inosine | 58-63-9 (Sigma-Aldrich) | 267.07349 | 5.0337 | 99.91 |
| Itaconate | Itaconic acid | 97-65-4 (Sigma-Aldrich) | 129.01931 | 2.4913 | 994.62 |
| L-Alanine | L-Alanine | 56-41-7 (Sigma-Aldrich) | 88.04042 | 8.5697 | 1003.48 |
| L-Arginine | L-Arginine | 74-79-3 (Sigma-Aldrich) | 173.10425 | 8.6333 | 1000.00 |
| L-Asparagine | L-Asparagine | 70-47-3 (Sigma-Aldrich) | 131.04613 | 8.2417 | 999.09 |
| L-Aspartate | L-Aspartic acid | 56-84-8 (Sigma-Aldrich) | 132.03016 | 8.5253 | 999.25 |
| L-Citrulline | Citrulline | 372-75-8 (Sigma-Aldrich) | 174.08826 | 8.3940 | 100.46 |
| L-Cysteic Acid | L-Cysteic acid monohydrate | 23537-25-9 (Sigma-Aldrich) | 167.99714 | 8.2720 | 999.09 |
| L-Cysteine | L-Cysteine hydrochloride | 52-89-1 (Sigma-Aldrich) | 120.01244 | 7.6363 | 1002.41 |
| L-Cystine | L-Cystine dihydrochloride | 30925-07-6 (Sigma-Aldrich) | 239.01655 | 8.7070 | 1001.21 |
| L-Dihydroorotate | L-Dihydroorotic acid | 5988-19-2 (Sigma-Aldrich) | 157.02546 | 5.5860 | 497.75 |
| L-Glutamate | L-Glutamic acid | 56-86-0 (Sigma-Aldrich) | 146.04584 | 8.3267 | 1004.55 |
| L-Glutamine | L-Glutamine | 56-85-9 (Sigma-Aldrich) | 145.06180 | 8.2327 | 1000.41 |
| L-Histidine | L-Histidine | 71-00-1 (Sigma-Aldrich) | 154.06225 | 8.6207 | 1000.32 |
| L-Isoleucine | L-Isoleucine | 73-32-5 (Sigma-Aldrich) | 130.08724 | 6.9747 | 1000.23 |
| L-Leucine | L-Leucine | 61-90-5 (Sigma-Aldrich) | 130.08723 | 6.8367 | 1013.95 |

| Chemical (pure compound) | Chemical Name | Manufacturer/ CAS No. | Average m/z | Average Retention Time (min) | Stock Concentration in Mixed Standard (μM) |
|---|---|---|---|---|---|
| L-Lysine | L-Lysine monohydrochloride | 657-27-2 (Sigma-Aldrich) | 145.09820 | 8.6703 | 1001.92 |
| L-Methionine | L-Methionine | 63-68-3 (Sigma-Aldrich) | 148.04378 | 7.1550 | 997.25 |
| L-Methionine Sulfoxide | L-Methionine sulfoxide | 3226-65-1 (Sigma-Aldrich) | 164.03873 | 8.3123 | 1001.15 |
| L-Ornithine | L-Ornithine monohydrochloride | 3184-13-2 (Sigma-Aldrich) | 131.08249 | 8.6353 | 97.85 |
| L-Phenylalanine | L-Phenylalanine | 63-91-2 (Sigma-Aldrich) | 164.07163 | 6.7800 | 1000.06 |
| L-Proline | L-Proline | 147-85-3 (Sigma-Aldrich) | 114.05602 | 7.6763 | 997.13 |
| L-Rhamnose | L-Rhamnose monohydrate | 10030-85-0 (Sigma-Aldrich) | 163.06123 | 2.0773 | 999.07 |
| L-Serine | L-Serine | 56-45-1 (Sigma-Aldrich) | 104.03533 | 8.2210 | 991.53 |
| L-Threonine | L-Threonine | 72-19-5 (Sigma-Aldrich) | 118.05097 | 8.0653 | 1000.67 |
| L-Tryptophan | L-Tryptophan | 73-22-3 (Sigma-Aldrich) | 203.08254 | 7.0403 | 998.87 |
| L-Tyrosine | L-Tyrosine | 60-18-4 (Sigma-Aldrich) | 180.06655 | 7.4307 | 1001.16 |
| L-Valine | L-Valine | 72-18-4 (Sigma-Aldrich) | 116.07169 | 7.4917 | 1003.84 |
| Myo-inositol | Myoinositol | 87-89-8 (Sigma-Aldrich) | 179.05603 | 6.9990 | 996.89 |
| N-Acetyl-Aspartate | N-acetyl-L-Aspartic acid | 997-55-7 (Sigma-Aldrich) | 174.04068 | 5.4640 | 998.06 |
| N-Acetyl-Glucosamine | N-acetyl-D-Glucosamine | 7512-17-6 (Sigma-Aldrich) | 220.08259 | 5.3530 | 1002.67 |
| N-Acetyl-Glutamate | N-acetyl-L-Glutamic acid | 1188-37-0 (Sigma-Aldrich) | 188.05639 | 4.9150 | 999.10 |
| N-Acetyl-Glutamine | Nα-acetyl-L-Glutamine | 2490-97-3 (Sigma-Aldrich) | 187.07233 | 5.8280 | 1000.11 |
| N-Acetyl-Glycine | N-acetyl-Glycine | 543-24-8 (Sigma-Aldrich) | 116.03530 | 3.7500 | 1005.98 |
| N-Acetyl-Leucine | N-acetyl-L-Leucine | 1188-21-2 (Sigma-Aldrich) | 172.09789 | 1.6423 | 999.94 |
| N-Acetyl-Methionine | N-Acetyl-Methionine | 1115-47-5 (Sigma-Aldrich) | 190.05425 | 2.6803 | 1000.78 |
| N-Acetyl-Phenylalanine | N-Acetyl-Phenylalanine | 2018-61-3 (Sigma-Aldrich) | 206.08215 | 2.4890 | 999.86 |
| Nicotinate | Nicotinic acid | 59-67-6 (Sigma-Aldrich) | 122.02472 | 3.0223 | 100.72 |
| O-Phospho-L-Serine | O-Phospho-L-Serine | 407-41-0 (Sigma-Aldrich) | 184.00152 | 9.0470 | 1002.86 |
| Orotate | Orotic acid | 65-86-1 (Sigma-Aldrich) | 155.00992 | 6.7730 | 99.94 |
| Pantothenate | D-Pantothenic acid hemicalcium salt | 137-08-6 (Sigma-Aldrich) | 218.10332 | 3.4907 | 99.89 |

| Chemical (pure compound) | Chemical Name | Manufacturer/ CAS No. | Average m/z | Average Retention Time (min) | Stock Concentration in Mixed Standard (μM) |
|---|---|---|---|---|---|
| Pyridoxine | Pyridoxine hydrochloride | 58-56-0 (Sigma-Aldrich) | 168.06658 | 6.4023 | 99.20 |
| Riboflavin | (-)-Riboflavin | 83-88-5 (Sigma-Aldrich) | 375.13112 | 4.6280 | 40.39 |
| Shikimate | Shikimic acid | 138-59-0 (Sigma-Aldrich) | 173.04540 | 3.6070 | 997.99 |
| Succinate | Sodium succinate dibasic hexahydrate | 6106-21-4 (Sigma-Aldrich) | 117.01931 | 1.7563 | 997.26 |
| Taurine | Taurine | 107-35-7 (Sigma-Aldrich) | 124.00730 | 7.0057 | 1005.19 |
| Thymidine | Thymidine | 50-89-5 (Sigma-Aldrich) | 241.08295 | 1.4193 | 99.08 |
| Thymine | Thymine | 65-71-4 (Sigma-Aldrich) | 125.03556 | 0.9683 | 99.12 |
| Uracil | Uracil | 66-22-8 (Sigma-Aldrich) | 111.01998 | 1.1543 | 99.03 |
| Uric Acid | Urate | 69-83-2 (Sigma-Aldrich) | 167.02103 | 5.6527 | 40.45 |
| Uridine | Uridine | 58-96-8 (Sigma-Aldrich) | 243.06216 | 3.1583 | 100.74 |
| Urocanate | Urocanic acid | 104-98-3 (Acros Organics) | 137.03559 | 4.4857 | 99.19 |
| Xanthine | Xanthine | 69-89-6 (Sigma-Aldrich) | 151.02618 | 3.7617 | 98.61 |
| Xanthosine | Xanthosine | 146-80-5 (Sigma-Aldrich) | 283.06840 | 5.1093 | 100.97 |

Table S2. Performance characteristics for visual inspection of standard curves, according to U.S. FDA Bioanalytical Method Validation Guidelines[1]*.

| **Compound** | **ULOQ (μM)** | **LLOQ (μM)** | **% of Standards within ±15% Nominal Concentration**** | **Intraday %RSD for Calculated Concentration (Minimum %)** | **Intraday %RSD for Calculated Concentration (Maximum %)** | **Quantitative (QUAN) or Qualitative (QUAL)?** |
|---|---|---|---|---|---|---|
| (R)-Mevalonate | 101.240 | 0.099 | 83% | 5.33% | 23.87% | QUAN |
| 2-Oxoglutarate | - | - | - | - | - | QUAL |
| Acetoacetate | 997.963 | 3.898 | 100% | 4.20% | 15.84% | QUAN |
| Acetyl-Threonine | 1002.639 | 0.979 | 83% | 1.21% | 20.65% | QUAN |
| Adenine | 25.346 | 0.025 | 83% | 0.91% | 12.51% | QUAN |
| Adenosine | - | - | - | - | - | QUAL |
| Adenosine 5'-monophosphate | 99.113 | 0.387 | 100% | 5.22% | 22.81% | QUAN |
| Biotin | 25.173 | 0.025 | 67% | 3.93% | 18.72% | QUAL |
| Carnosine | - | - | - | - | - | QUAL |
| Creatinine | - | - | - | - | - | QUAL |
| Cytidine | 98.676 | 0.096 | 100% | 6.37% | 16.26% | QUAN |
| D-Arabitol | 249.096 | 0.973 | 80% | 5.08% | 20.38% | QUAN |
| D-Fructose 1,6-bisphosphate | 999.852 | 3.906 | 60% | 2.34% | 23.88% | QUAL |
| D-Glucose | 249.500 | 0.975 | 100% | 12.30% | 38.57% | QUAN |
| D-Glucose 6-phosphate | 998.157 | 0.975 | 100% | 3.49% | 21.39% | QUAN |
| DL-Homocysteine | - | - | - | - | - | QUAL |
| Folate | - | - | - | - | - | QUAL |
| Fumarate | 1001.000 | 0.244 | 71% | 5.40% | 29.55% | QUAL |
| Glycine | - | - | - | - | - | QUAL |
| Guanine | 24.978 | 0.098 | 67% | 6.19% | 24.03% | QUAL |
| Guanosine | 100.268 | 0.392 | 100% | 4.52% | 20.98% | QUAN |
| Hippurate | 15.675 | 0.015 | 83% | 1.82% | 18.99% | QUAN |
| Hypoxanthine | 24.429 | 0.024 | 50% | 1.53% | 21.44% | QUAL |
| Inosine | 24.979 | 0.098 | 80% | 5.58% | 21.84% | QUAN |

| Compound | ULOQ (μM) | LLOQ (μM) | % of Standards within ±15% Nominal Concentration** | Intraday %RSD for Calculated Concentration (Minimum %) | Intraday %RSD for Calculated Concentration (Maximum %) | Quantitative (QUAN) or Qualitative (QUAL)? |
|---|---|---|---|---|---|---|
| Itaconate | 62.164 | 0.061 | 83% | 2.32% | 17.09% | QUAN |
| L-Alanine | - | - | - | - | - | QUAL |
| L-Arginine | 1000.000 | 0.977 | 67% | 2.44% | 13.07% | QUAL |
| L-Asparagine | 999.092 | 0.976 | 67% | 3.61% | 13.87% | QUAL |
| L-Aspartate | 999.249 | 3.903 | 80% | 1.30% | 24.78% | QUAN |
| L-Citrulline | 100.462 | 0.392 | 80% | 1.18% | 20.03% | QUAN |
| L-Cysteate | 249.773 | 0.244 | 67% | 1.18% | 28.06% | QUAL |
| L-Cysteine | - | - | - | - | - | QUAL |
| L-Cystine | 1001.213 | 3.911 | 83% | 7.00% | 16.89% | QUAN |
| L-Dihydroorotate | 497.755 | 0.122 | 100% | 6.26% | 18.84% | QUAN |
| L-Glutamate | - | - | - | - | - | QUAL |
| L-Glutamine | 250.103 | 0.977 | 100% | 2.75% | 13.32% | QUAN |
| L-Histidine | 1000.322 | 0.977 | 83% | 4.67% | 20.14% | QUAN |
| L-Isoleucine | - | - | - | - | - | QUAL |
| L-Leucine | - | - | - | - | - | QUAL |
| L-Lysine | - | - | - | - | - | QUAL |
| L-Methionine | 997.252 | 3.896 | 80% | 1.37% | 17.74% | QUAN |
| L-Methionine Sulfoxide | 1001.150 | 0.978 | 67% | 7.12% | 15.45% | QUAL |
| L-Ornithine | 97.853 | 0.382 | 80% | 3.25% | 18.78% | QUAN |
| L-Phenylalanine | - | - | - | - | - | QUAL |
| L-Proline | - | - | - | - | - | QUAL |
| L-Rhamnose | - | - | - | - | - | QUAL |
| L-Serine | 247.883 | 0.968 | 60% | 2.37% | 21.81% | QUAL |
| L-Threonine | 1000.672 | 3.909 | 100% | 2.34% | 8.74% | QUAN |
| L-Tryptophan | 998.874 | 3.902 | 60% | 2.78% | 26.63% | QUAL |
| L-Tyrosine | 1001.159 | 0.978 | 83% | 3.30% | 21.95% | QUAN |

| Compound | ULOQ (μM) | LLOQ (μM) | % of Standards within ±15% Nominal Concentration** | Intraday %RSD for Calculated Concentration (Minimum %) | Intraday %RSD for Calculated Concentration (Maximum %) | Quantitative (QUAN) or Qualitative (QUAL)? |
|---|---|---|---|---|---|---|
| L-Valine | - | - | - | - | - | QUAL |
| myo-Inositol | - | - | - | - | - | QUAL |
| N-Acetyl-D-glucosamine | 250.667 | 0.979 | 100% | 4.59% | 14.44% | QUAN |
| N-Acetyl-glycine | 251.494 | 0.246 | 57% | 6.10% | 25.97% | QUAL |
| N-Acetyl-L-aspartate | 249.515 | 0.975 | 100% | 2.86% | 12.84% | QUAN |
| N-Acetyl-L-glutamate | 999.101 | 0.244 | 100% | 5.70% | 21.71% | QUAN |
| N-Acetyl-L-Leucine | 62.496 | 0.061 | 100% | 1.93% | 16.77% | QUAN |
| N-Acetyl-L-methionine | 62.549 | 0.244 | 100% | 6.43% | 12.00% | QUAN |
| N-Acetyl-L-phenylalanine | 249.964 | 0.976 | 80% | 1.35% | 14.38% | QUAN |
| N-alpha-Acetyl-L-glutamine | 1000.106 | 0.244 | 86% | 4.07% | 17.92% | QUAN |
| Nicotinate | 6.295 | 0.025 | 100% | 2.60% | 30.85% | QUAN |
| O-Phospho-L-serine | 1002.864 | 3.917 | 80% | 6.99% | 21.50% | QUAN |
| Orotate | 99.936 | 0.390 | 80% | 2.19% | 9.08% | QUAN |
| Pantothenate | 99.887 | 0.024 | 100% | 8.85% | 28.85% | QUAN |
| Pyridoxine | 99.202 | 0.097 | 67% | 1.27% | 17.65% | QUAL |
| Riboflavin | 40.387 | 0.631 | 75% | 15.99% | 29.02% | QUAN |
| Shikimate | 249.498 | 0.244 | 83% | 5.29% | 31.99% | QUAN |
| Succinate | 62.329 | 0.243 | 100% | 3.62% | 20.10% | QUAN |
| Taurine | 251.298 | 0.245 | 83% | 2.14% | 12.45% | QUAN |
| Thymidine | 24.770 | 0.097 | 100% | 3.79% | 29.35% | QUAN |
| Thymine | 99.120 | 0.387 | 80% | 17.18% | 28.81% | QUAN |
| Uracil | 24.757 | 0.097 | 100% | 3.85% | 8.79% | QUAN |
| Urate | 40.450 | 0.040 | 83% | 5.77% | 31.89% | QUAN |
| Uridine | 25.184 | 0.025 | 50% | 4.35% | 21.03% | QUAL |
| Urocanate | 6.199 | 0.024 | 100% | 5.16% | 13.37% | QUAN |
| Xanthine | 98.613 | 0.024 | 86% | 2.15% | 15.39% | QUAN |

| Compound | ULOQ (μM) | LLOQ (μM) | % of Standards within ±15% Nominal Concentration** | Intraday %RSD for Calculated Concentration (Minimum %) | Intraday %RSD for Calculated Concentration (Maximum %) | Quantitative (QUAN) or Qualitative (QUAL)? |
|---|---|---|---|---|---|---|
| Xanthosine | 100.975 | 0.099 | 50% | 7.46% | 17.13% | QUAL |

*Performance characteristics evaluated include the upper limit of quantification (ULOQ) – the highest amount of an analyte in a sample that can be determined with precision and accuracy and the lower limit of quantification (LLOQ) – the lowest amount of an analyte that can be quantitatively determined with acceptable precision and accuracy. The U.S. FDA guidelines also state that non-zero calibrators should be within ±15% of the nominal (expected) concentration, except at the LLOQ where the calibrators should be ±20% of the nominal concentration in each validation run and that 75% and a minimum of six non-zero calibrator levels should meet these criteria in each validation run. We used these criteria, along with the intraday relative standard deviation (%RSD) in calculated concentration between 3 technical replicates, to determine the linear range for each compound according to visual inspection of standard curves. We considered that compounds that did not meet these criteria be used for qualitative purposes only and we did not show data for compounds that had shown no correlation between signal intensity and concentration.

**Calibrator levels at the LLOQ were considered acceptable if the calculated concentration was within ±20% of the nominal concentration.

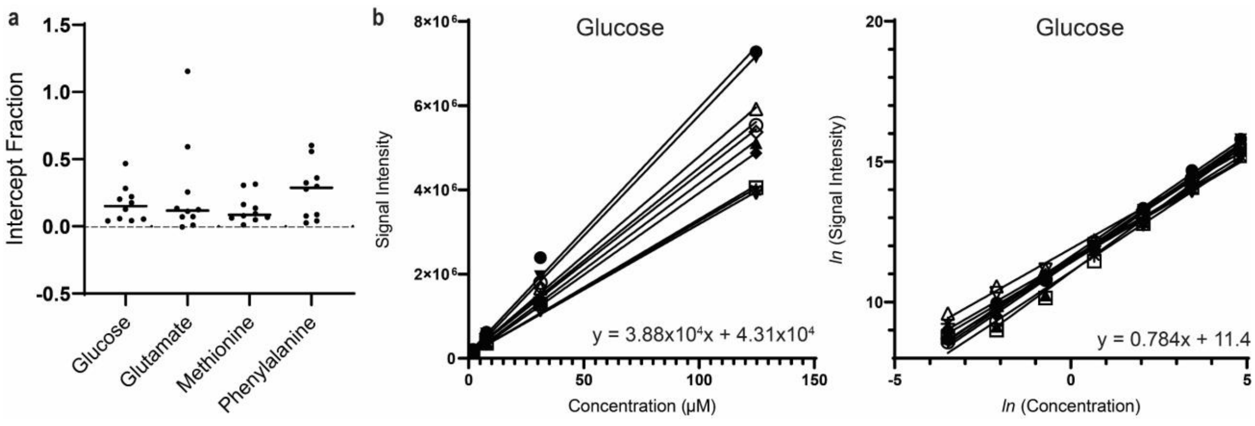


Figure S1. (a) Examples of intercept fraction values ($b/(m \times \overline{x}\,)$) for the standard curves of ten metabolites run over ten batches. (b) Examples of a set of ten standard curves for glucose in the linear and natural logarithmic scale. Each curve corresponds to one batch and the line of best fit equations show the slope and intercept from one standard curve.

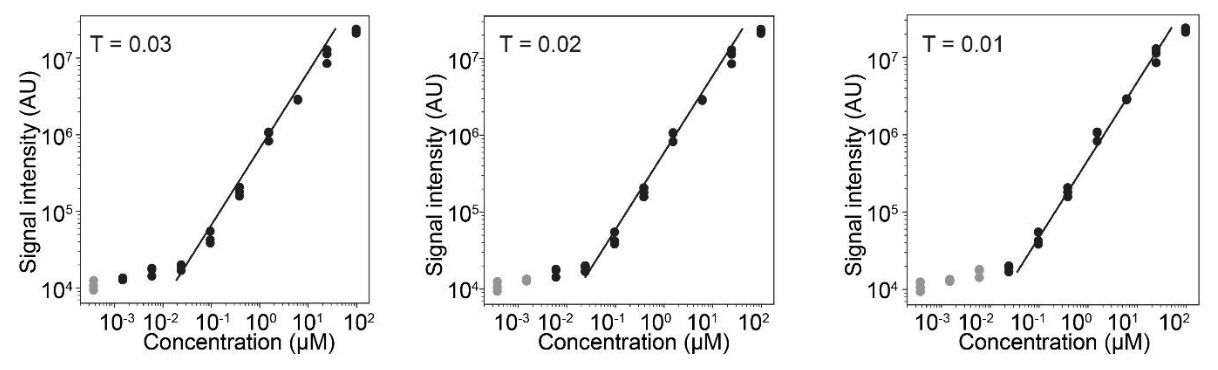


Figure S2. Effect of using different threshold residual values (T) to stop the algorithm's fitting process for hypoxanthine, using a 10-point standard curve with 3 technical replicates. Black dots show points belonging to the linear range while gray dots show points outside the linear range.

Table S3. SCALiR performance evaluation results for 10 representative compounds, showing the concentrations calculated by the SCALiR app compared to the expected concentration in an 8-point standard curve. The average concentration (µM) of each standard across 10 analytical batches and coefficient of variation (CV) are shown for each standard, as well as performance evaluation according to the U.S. FDA guidelines (non-zero calibrators should be within ±15% of the nominal (expected) concentration)[1].

| **Compound/ Standard Number** | **Expected Concentration (µM)** | **SCALiR-Calculated Concentration (µM)** | | | | | | | | | | **Average Calculated Concentration (µM)** | **CV** | **Within ± 15% Expected Concentration?** |
|---|---|---|---|---|---|---|---|---|---|---|---|---|---|---|
| | | Batch 1 | Batch 2 | Batch 3 | Batch 4 | Batch 5 | Batch 6 | Batch 7 | Batch 8 | Batch 9 | Batch 10 | | | |
| Acetyl-threonine | | | | | | | | | | | | | | |
| Std 1 | 0.031 | 0.044 | 0.042 | 0.041 | 0.041 | 0.034 | 0.046 | 0.058 | 0.037 | 0.036 | 0.063 | 0.044 | 0.210 | FALSE |
| Std 2 | 0.122 | 0.146 | 0.126 | 0.110 | 0.115 | 0.133 | 0.124 | 0.143 | 0.133 | 0.135 | 0.132 | 0.130 | 0.087 | TRUE |
| Std 3 | 0.490 | 0.504 | 0.488 | 0.553 | 0.549 | 0.642 | 0.369 | 0.597 | 0.551 | 0.559 | 0.582 | 0.539 | 0.137 | TRUE |
| Std 4 | 1.958 | 1.523 | 2.189 | 1.701 | 2.012 | 2.075 | 1.839 | 1.972 | 2.115 | 2.081 | 2.129 | 1.964 | 0.108 | TRUE |
| Std 5 | 7.834 | 8.210 | 8.700 | 9.875 | 8.406 | 6.960 | 7.932 | 8.241 | 8.523 | 10.002 | 8.284 | 8.513 | 0.104 | TRUE |
| Std 6 | 31.335 | 28.376 | 30.199 | 39.280 | 28.400 | 37.419 | 41.678 | 27.983 | 34.840 | 34.281 | 36.271 | 33.873 | 0.145 | TRUE |
| Std 7 | 125.341 | 96.726 | 117.201 | 108.499 | 125.688 | 117.028 | 124.783 | 93.043 | 104.072 | 107.479 | 108.797 | 110.332 | 0.099 | TRUE |
| Std 8 | 501.365 | 248.631 | 314.931 | 312.537 | 352.805 | 298.880 | 344.424 | 193.641 | 313.882 | 267.045 | 339.079 | 298.585 | 0.165 | FALSE |
| Adenine | | | | | | | | | | | | | | |
| Std 1 | 0.003 | 0.006 | 0.005 | 0.005 | 0.009 | 0.004 | 0.009 | 0.007 | 0.006 | 0.006 | 0.010 | 0.006 | 0.007 | FALSE |
| Std 2 | 0.012 | 0.014 | 0.014 | 0.014 | 0.019 | 0.015 | 0.018 | 0.016 | 0.016 | 0.013 | 0.014 | 0.014 | 0.015 | FALSE |
| Std 3 | 0.050 | 0.059 | 0.046 | 0.048 | 0.061 | 0.058 | 0.052 | 0.056 | 0.059 | 0.060 | 0.063 | 0.059 | 0.056 | TRUE |
| Std 4 | 0.198 | 0.214 | 0.195 | 0.180 | 0.215 | 0.188 | 0.206 | 0.210 | 0.189 | 0.228 | 0.182 | 0.214 | 0.201 | TRUE |
| Std 5 | 0.792 | 0.826 | 0.677 | 0.663 | 0.754 | 0.711 | 0.794 | 0.869 | 0.729 | 0.970 | 0.716 | 0.826 | 0.771 | TRUE |
| Std 6 | 3.168 | 2.925 | 2.376 | 2.501 | 2.469 | 2.257 | 2.985 | 2.882 | 2.335 | 2.764 | 2.601 | 2.925 | 2.609 | FALSE |
| Std 7 | 12.673 | 9.073 | 6.640 | 6.515 | 6.936 | 6.296 | 8.252 | 8.166 | 6.711 | 7.841 | 6.441 | 9.073 | 7.287 | FALSE |
| Std 8 | 50.692 | 20.705 | 16.825 | 14.866 | 17.314 | 16.477 | 18.946 | 20.073 | 15.956 | 19.569 | 14.133 | 20.705 | 17.487 | FALSE |
| Asparagine | | | | | | | | | | | | | | |
| Std 1 | 0.030 | 0.104 | 0.148 | 0.080 | 0.111 | 0.075 | 0.079 | 0.143 | 0.126 | 0.058 | 0.110 | 0.049 | 1.346 | FALSE |
| Std 2 | 0.122 | 0.166 | 0.241 | 0.145 | 0.213 | 0.142 | 0.173 | 0.159 | 0.175 | 0.129 | 0.178 | 0.172 | 0.246 | FALSE |
| Std 3 | 0.488 | 0.513 | 0.623 | 0.519 | 0.660 | 0.534 | 0.500 | 0.552 | 0.511 | 0.496 | 0.617 | 0.521 | 0.185 | TRUE |
| Std 4 | 1.951 | 1.840 | 2.183 | 1.831 | 2.261 | 1.962 | 2.050 | 1.781 | 2.056 | 1.727 | 1.889 | 1.673 | 0.102 | TRUE |
| Std 5 | 7.805 | 7.447 | 8.606 | 7.774 | 8.730 | 7.312 | 7.824 | 7.852 | 8.496 | 7.188 | 7.495 | 7.085 | 0.113 | TRUE |
| Std 6 | 31.222 | 28.456 | 31.205 | 30.214 | 29.811 | 31.898 | 31.106 | 30.268 | 30.310 | 28.635 | 29.298 | 31.560 | 0.208 | TRUE |
| Std 7 | 124.886 | 113.179 | 104.579 | 119.120 | 96.674 | 118.627 | 104.572 | 123.358 | 104.779 | 111.747 | 95.176 | 129.101 | 0.142 | TRUE |
| Std 8 | 499.546 | 476.989 | 383.142 | 464.473 | 389.970 | 433.760 | 396.626 | 390.250 | 361.838 | 368.691 | 411.550 | 474.016 | 0.093 | TRUE |
| Fumarate | | | | | | | | | | | | | | |

| Compound/ Standard Number | Expected Concentration (µM) | SCALiR-Calculated Concentration (µM) | | | | | | | | | | Average Calculated Concentration (µM) | CV | Within ± 15% Expected Concentration? |
|---|---|---|---|---|---|---|---|---|---|---|---|---|---|---|
| | | Batch 1 | Batch 2 | Batch 3 | Batch 4 | Batch 5 | Batch 6 | Batch 7 | Batch 8 | Batch 9 | Batch 10 | | | |
| Std 1 | 0.031 | 0.104 | 0.148 | 0.080 | 0.111 | 0.075 | 0.079 | 0.143 | 0.126 | 0.058 | 0.110 | 0.104 | 0.291 | FALSE |
| Std 2 | 0.122 | 0.166 | 0.241 | 0.145 | 0.213 | 0.142 | 0.173 | 0.159 | 0.175 | 0.129 | 0.178 | 0.172 | 0.196 | FALSE |
| Std 3 | 0.489 | 0.513 | 0.623 | 0.519 | 0.660 | 0.534 | 0.500 | 0.552 | 0.511 | 0.496 | 0.617 | 0.553 | 0.107 | TRUE |
| Std 4 | 1.955 | 1.840 | 2.183 | 1.831 | 2.261 | 1.962 | 2.050 | 1.781 | 2.056 | 1.727 | 1.889 | 1.958 | 0.090 | TRUE |
| Std 5 | 7.820 | 7.447 | 8.606 | 7.774 | 8.730 | 7.312 | 7.824 | 7.852 | 8.496 | 7.188 | 7.495 | 7.872 | 0.071 | TRUE |
| Std 6 | 31.281 | 28.456 | 31.205 | 30.214 | 29.811 | 31.898 | 31.106 | 30.268 | 30.310 | 28.635 | 29.298 | 30.120 | 0.037 | TRUE |
| Std 7 | 125.125 | 113.179 | 104.579 | 119.120 | 96.674 | 118.627 | 104.572 | 123.358 | 104.779 | 111.747 | 95.176 | 109.181 | 0.088 | TRUE |
| Std 8 | 500.500 | 476.989 | 383.142 | 464.473 | 389.970 | 433.760 | 396.626 | 390.250 | 361.838 | 368.691 | 411.550 | 407.729 | 0.096 | FALSE |
| Hypoxanthine | | | | | | | | | | | | | | |
| Std 1 | 0.003 | 0.006 | 0.008 | 0.005 | 0.009 | 0.005 | 0.010 | 0.007 | 0.006 | 0.006 | 0.010 | 0.007 | 0.263 | FALSE |
| Std 2 | 0.012 | 0.018 | 0.020 | 0.014 | 0.025 | 0.014 | 0.023 | 0.014 | 0.017 | 0.015 | 0.014 | 0.017 | 0.232 | FALSE |
| Std 3 | 0.048 | 0.051 | 0.057 | 0.043 | 0.064 | 0.048 | 0.062 | 0.049 | 0.051 | 0.055 | 0.061 | 0.054 | 0.125 | TRUE |
| Std 4 | 0.191 | 0.211 | 0.215 | 0.194 | 0.206 | 0.174 | 0.215 | 0.218 | 0.176 | 0.202 | 0.200 | 0.201 | 0.078 | TRUE |
| Std 5 | 0.763 | 0.777 | 0.798 | 0.800 | 0.788 | 0.741 | 0.781 | 0.849 | 0.674 | 0.773 | 0.820 | 0.780 | 0.060 | TRUE |
| Std 6 | 3.054 | 3.200 | 2.989 | 2.916 | 2.823 | 2.704 | 3.049 | 3.508 | 2.512 | 2.675 | 2.817 | 2.919 | 0.098 | TRUE |
| Std 7 | 12.214 | 10.578 | 8.932 | 10.958 | 8.882 | 8.514 | 8.163 | 10.718 | 7.125 | 8.920 | 8.036 | 9.083 | 0.140 | FALSE |
| Std 8 | 48.858 | 30.397 | 20.585 | 29.621 | 22.008 | 24.611 | 22.670 | 31.287 | 17.568 | 26.493 | 18.742 | 24.398 | 0.201 | FALSE |
| Hippurate | | | | | | | | | | | | | | |
| Std 1 | 0.031 | 0.037 | 0.032 | 0.034 | 0.049 | 0.036 | 0.054 | 0.038 | 0.034 | 0.033 | 0.063 | 0.041 | 0.257 | FALSE |
| Std 2 | 0.122 | 0.123 | 0.136 | 0.139 | 0.137 | 0.128 | 0.132 | 0.126 | 0.129 | 0.135 | 0.130 | 0.132 | 0.039 | TRUE |
| Std 3 | 0.490 | 0.547 | 0.530 | 0.475 | 0.545 | 0.554 | 0.549 | 0.499 | 0.519 | 0.524 | 0.593 | 0.533 | 0.061 | TRUE |
| Std 4 | 1.959 | 1.912 | 1.934 | 1.872 | 2.039 | 1.933 | 2.081 | 1.675 | 2.029 | 2.005 | 1.839 | 1.932 | 0.061 | TRUE |
| Std 5 | 7.838 | 5.966 | 6.341 | 6.708 | 6.055 | 5.632 | 6.099 | 7.052 | 6.123 | 6.037 | 6.494 | 6.251 | 0.065 | FALSE |
| Std 6 | 31.351 | 14.467 | 15.058 | 13.524 | 15.378 | 13.499 | 16.170 | 13.269 | 16.262 | 14.202 | 13.889 | 14.572 | 0.075 | FALSE |
| Std 7 | 125.404 | 27.039 | 33.120 | 24.646 | 32.553 | 24.841 | 33.055 | 30.884 | 36.171 | 24.068 | 27.722 | 29.410 | 0.146 | FALSE |
| Std 8 | 501.616 | 49.032 | 59.598 | 45.326 | 64.319 | 45.912 | 48.676 | 51.794 | 62.110 | 42.521 | 49.741 | 51.903 | 0.145 | FALSE |
| N-acetyl-Leucine | | | | | | | | | | | | | | |
| Std 1 | 0.031 | 0.037 | 0.034 | 0.034 | 0.046 | 0.038 | 0.055 | 0.036 | 0.034 | 0.033 | 0.062 | 0.041 | 0.242 | FALSE |
| Std 2 | 0.122 | 0.127 | 0.133 | 0.143 | 0.134 | 0.125 | 0.144 | 0.128 | 0.122 | 0.134 | 0.153 | 0.134 | 0.073 | TRUE |
| Std 3 | 0.488 | 0.534 | 0.577 | 0.507 | 0.506 | 0.543 | 0.529 | 0.492 | 0.581 | 0.597 | 0.562 | 0.543 | 0.065 | TRUE |
| Std 4 | 1.953 | 2.090 | 1.952 | 1.937 | 1.847 | 2.233 | 2.072 | 2.005 | 2.003 | 2.066 | 2.056 | 2.026 | 0.052 | TRUE |
| Std 5 | 7.812 | 7.620 | 7.409 | 7.370 | 7.046 | 6.990 | 7.948 | 8.138 | 8.058 | 7.404 | 7.302 | 7.528 | 0.054 | TRUE |

| Compound/ Standard Number | Expected Concentration (µM) | SCALiR-Calculated Concentration (µM) | | | | | | | | | | Average Calculated Concentration (µM) | CV | Within ± 15% Expected Concentration? |
|---|---|---|---|---|---|---|---|---|---|---|---|---|---|---|
| | | Batch 1 | Batch 2 | Batch 3 | Batch 4 | Batch 5 | Batch 6 | Batch 7 | Batch 8 | Batch 9 | Batch 10 | | | |
| Std 6 | 31.248 | 21.478 | 22.866 | 24.343 | 21.147 | 21.637 | 22.619 | 23.255 | 21.995 | 21.173 | 21.979 | 22.249 | 0.046 | FALSE |
| Std 7 | 124.993 | 55.685 | 62.124 | 56.169 | 48.477 | 49.362 | 56.609 | 51.866 | 61.478 | 45.269 | 50.025 | 53.706 | 0.105 | FALSE |
| Std 8 | 499.971 | 100.431 | 111.681 | 119.268 | 101.237 | 97.607 | 121.387 | 114.329 | 125.366 | 97.073 | 104.416 | 109.280 | 0.096 | FALSE |
| Nicotinate | | | | | | | | | | | | | | |
| Std 1 | 0.003 | 0.005 | 0.008 | 0.004 | 0.011 | 0.007 | 0.010 | 0.008 | 0.008 | 0.007 | 0.010 | 0.008 | 0.264 | FALSE |
| Std 2 | 0.012 | 0.013 | 0.023 | 0.016 | 0.023 | 0.015 | 0.023 | 0.016 | 0.021 | 0.015 | 0.022 | 0.019 | 0.217 | FALSE |
| Std 3 | 0.049 | 0.047 | 0.052 | 0.053 | 0.064 | 0.063 | 0.059 | 0.055 | 0.062 | 0.054 | 0.060 | 0.057 | 0.096 | FALSE |
| Std 4 | 0.197 | 0.198 | 0.242 | 0.193 | 0.232 | 0.234 | 0.224 | 0.203 | 0.223 | 0.238 | 0.202 | 0.219 | 0.083 | TRUE |
| Std 5 | 0.787 | 0.778 | 0.830 | 0.685 | 0.829 | 0.893 | 0.866 | 0.785 | 0.831 | 0.852 | 0.770 | 0.812 | 0.073 | TRUE |
| Std 6 | 3.148 | 2.917 | 2.809 | 2.821 | 2.820 | 3.124 | 2.774 | 2.873 | 2.858 | 2.897 | 2.593 | 2.848 | 0.046 | TRUE |
| Std 7 | 12.590 | 8.198 | 10.259 | 8.377 | 8.683 | 10.140 | 9.551 | 9.136 | 9.196 | 7.862 | 7.866 | 8.927 | 0.098 | FALSE |
| Std 8 | 50.361 | 23.731 | 26.400 | 22.877 | 24.302 | 30.272 | 21.294 | 23.364 | 30.175 | 20.740 | 19.639 | 24.279 | 0.151 | FALSE |
| Pantothenate | | | | | | | | | | | | | | |
| Std 1 | 0.003 | 0.004 | 0.006 | 0.004 | 0.007 | 0.004 | 0.007 | 0.004 | 0.005 | 0.004 | 0.008 | 0.005 | 0.277 | FALSE |
| Std 2 | 0.012 | 0.013 | 0.015 | 0.015 | 0.013 | 0.016 | 0.017 | 0.014 | 0.014 | 0.013 | 0.015 | 0.015 | 0.091 | FALSE |
| Std 3 | 0.049 | 0.057 | 0.053 | 0.052 | 0.061 | 0.058 | 0.052 | 0.051 | 0.049 | 0.050 | 0.066 | 0.055 | 0.098 | TRUE |
| Std 4 | 0.195 | 0.218 | 0.224 | 0.191 | 0.207 | 0.201 | 0.220 | 0.225 | 0.188 | 0.196 | 0.206 | 0.208 | 0.065 | TRUE |
| Std 5 | 0.780 | 0.739 | 0.789 | 0.785 | 0.771 | 0.643 | 0.803 | 0.852 | 0.755 | 0.661 | 0.765 | 0.756 | 0.083 | TRUE |
| Std 6 | 3.121 | 2.602 | 2.778 | 2.729 | 2.178 | 2.800 | 2.721 | 2.556 | 2.792 | 2.295 | 2.873 | 2.632 | 0.087 | FALSE |
| Std 7 | 12.486 | 8.371 | 9.126 | 7.913 | 7.819 | 8.393 | 8.291 | 7.720 | 8.654 | 7.022 | 8.072 | 8.138 | 0.070 | FALSE |
| Std 8 | 49.943 | 20.941 | 21.309 | 27.250 | 20.236 | 20.077 | 21.702 | 21.770 | 21.739 | 18.527 | 21.007 | 21.456 | 0.106 | FALSE |
| Taurine | | | | | | | | | | | | | | |
| Std 1 | 0.031 | 0.025 | 0.043 | 0.041 | 0.050 | 0.040 | 0.096 | 0.048 | 0.034 | 0.035 | 0.093 | 0.051 | 0.479 | FALSE |
| Std 2 | 0.123 | 0.156 | 0.122 | 0.131 | 0.148 | 0.130 | 0.153 | 0.122 | 0.127 | 0.157 | 0.131 | 0.138 | 0.104 | TRUE |
| Std 3 | 0.491 | 0.541 | 0.524 | 0.492 | 0.513 | 0.477 | 0.483 | 0.545 | 0.578 | 0.530 | 0.601 | 0.528 | 0.075 | TRUE |
| Std 4 | 1.963 | 2.239 | 1.820 | 2.116 | 1.670 | 2.130 | 1.792 | 2.189 | 2.030 | 2.197 | 1.820 | 2.000 | 0.103 | TRUE |
| Std 5 | 7.853 | 8.871 | 7.264 | 8.410 | 7.622 | 8.611 | 7.989 | 7.477 | 7.543 | 7.847 | 7.769 | 7.940 | 0.066 | TRUE |
| Std 6 | 31.412 | 31.377 | 31.994 | 34.634 | 28.571 | 27.695 | 35.688 | 28.319 | 32.722 | 31.113 | 36.853 | 31.897 | 0.099 | TRUE |
| Std 7 | 125.649 | 120.874 | 123.790 | 108.281 | 110.544 | 129.289 | 137.315 | 120.796 | 125.264 | 121.076 | 121.143 | 121.837 | 0.068 | TRUE |
| Std 8 | 502.597 | 354.635 | 395.042 | 317.705 | 371.979 | 344.825 | 355.264 | 316.597 | 360.082 | 300.052 | 370.970 | 348.715 | 0.084 | FALSE |

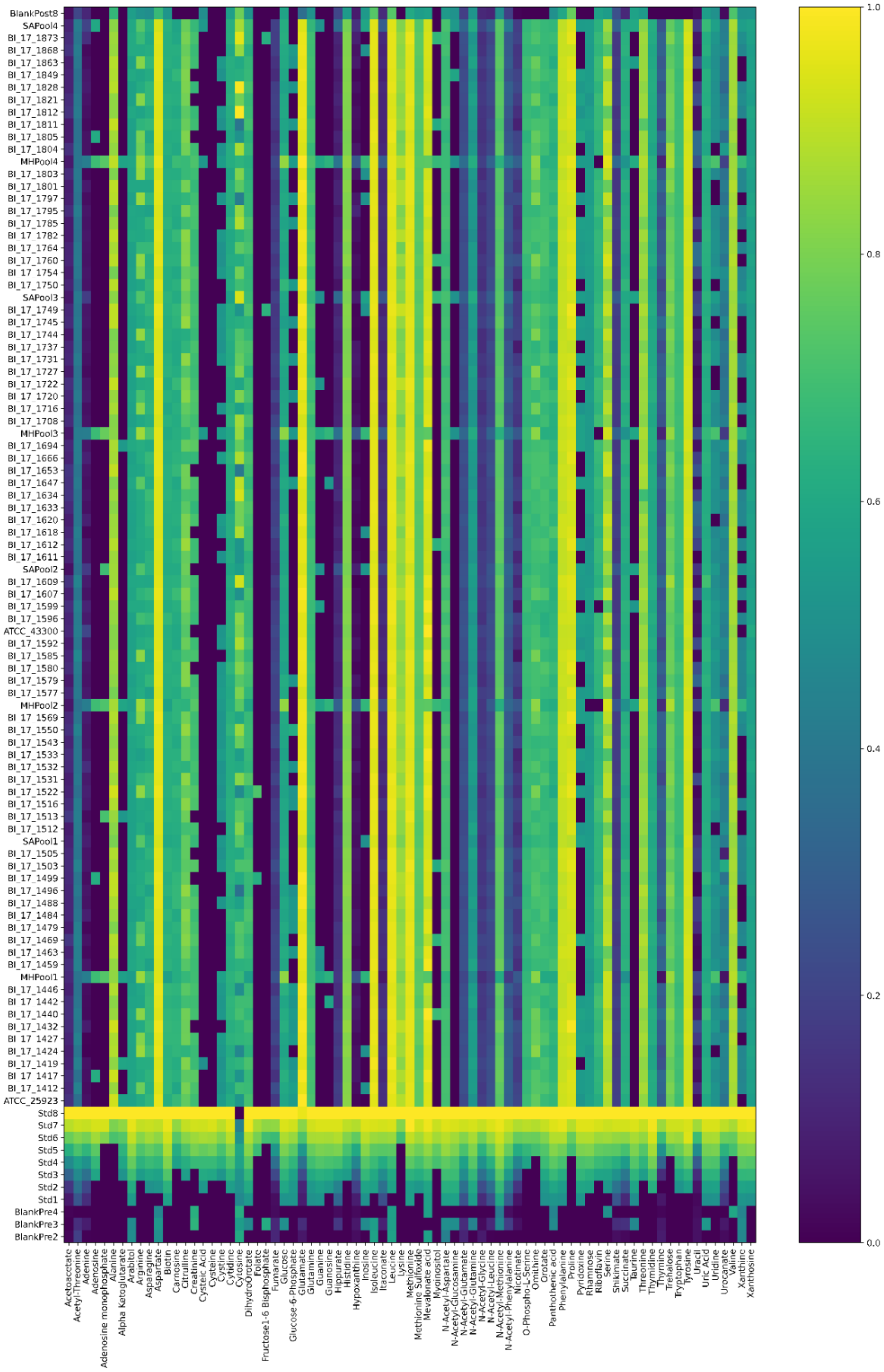


Figure S3. Heatmap of bacterial metabolomics dataset showing normalized metabolite concentrations in one batch of samples.

### References

(1) U.S. FDA. *Guidance for Industry Bioanalytical Method Validation Guidance for Industry Bioanalytical Method Validation*; 2018.
