## Supplementary material for "SCALiR: a web application for automating absolute quantification of mass spectrometry-based metabolomics data": Tutorial

**SCALiR Tutorial**

This quick (~5-10 minute) tutorial helps users understand different features of SCALiR and how to visualize and customize data generated by the app. Instructions are also available on the main page of the app.

**Step 1.** Create a new folder on your device to store the sample data upload files. Open the SCALiR web application and on the left-hand side of the screen, click on the link to the standards concentration file. Repeat this step for the three peaklist data files (two for Mint data and one for Maven data).

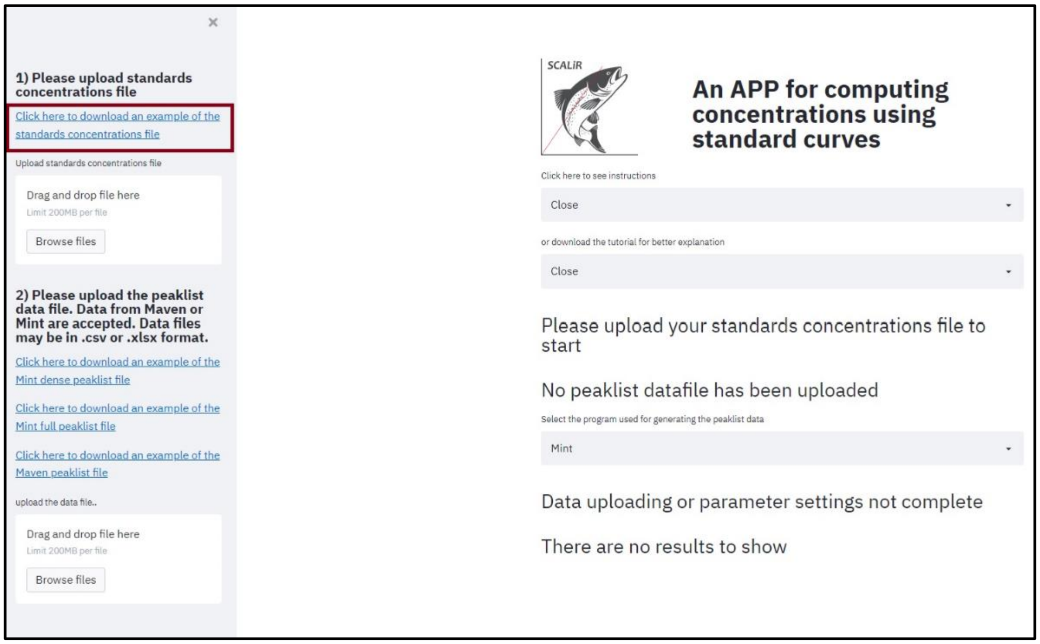

**Step 2.** On the web application, upload the standards concentration file and the "SCALiR_Maven_Peaklist_sample" file. The data will now pop up on the screen. Scroll down and select "Maven" in the program selection menu. The default option for slope fitting is "Fixed fit" and you should see the standard curve parameters and concentration datafiles. You can click on the highlighted links to download these files.

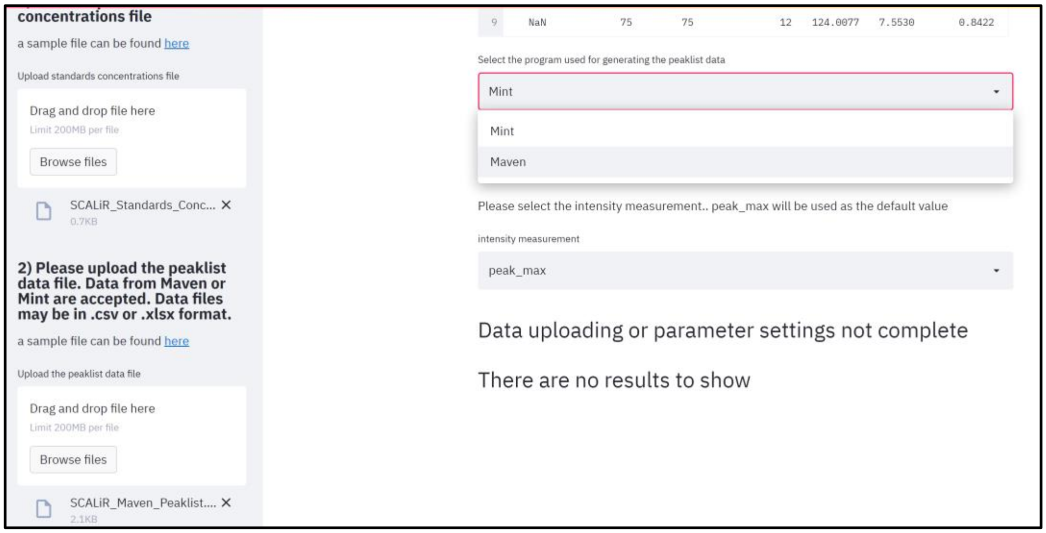


**Step 3.** Scroll down to the bottom of the page. The app is displaying the default metabolite. Click on the compound menu and select "Fumarate". You can now change the axis labels on the displayed graph.

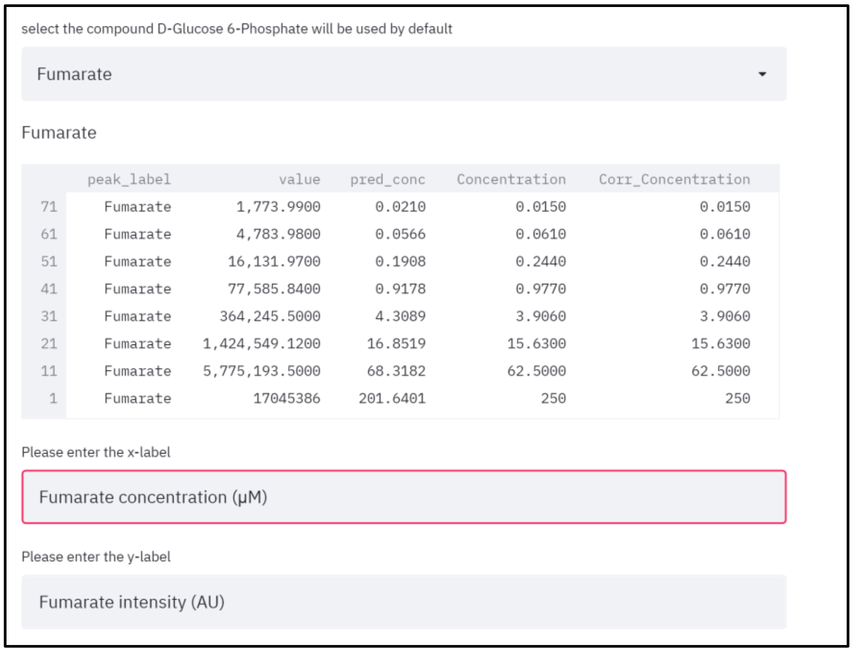

**Step 4.** Change the x-axis label to “Fumarate concentration (mM)” and click enter. Then change the y-axis label to “Fumarate peak area (AU)” and click enter. If you would like to leave the axis labels blank, delete all text in the box and click enter. To download the plot, right-click and select “Save as.”.

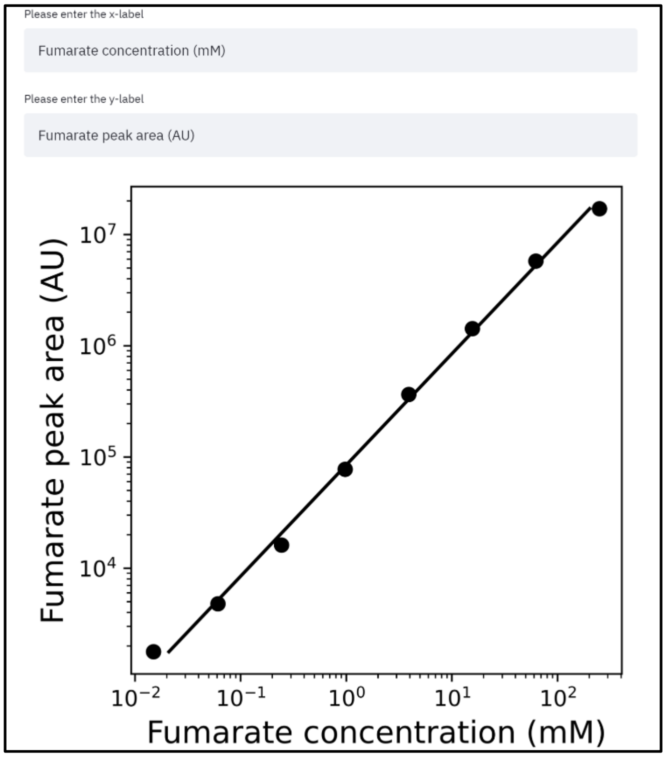


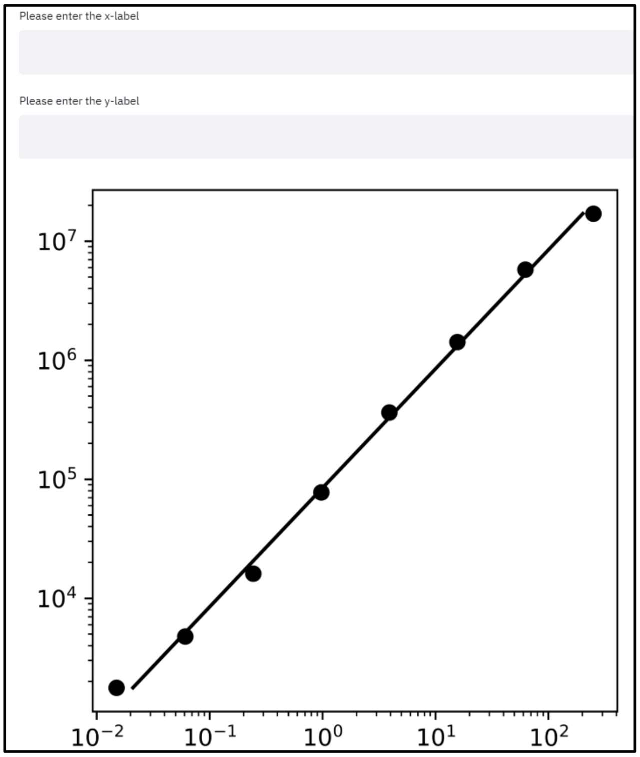

**Step 5.** Refresh the browser. Upload the sample standards concentrations and the "SCALiR_MINT Peaklist_Full_Results_sample" files. The default program and table type are Mint and full results, so you do not need to change those but can select "peak_area" under the intensity measurement menu. Also, select "Interval fit" under the line of best fit menu. Change the slider bar to 0.90 – 1.10 from the recommended interval of 0.85 to 1.15. SCALiR will re-run the algorithm and your new results will pop up below as before so you can download the data files as well as standard curve visualizations. To use the "SCALiR_MINT_Peaklist_Dense_Peak_Max_sample" file, select "dense peak_max" for the type of table used (peak max is the only intensity measurement available for this file type).

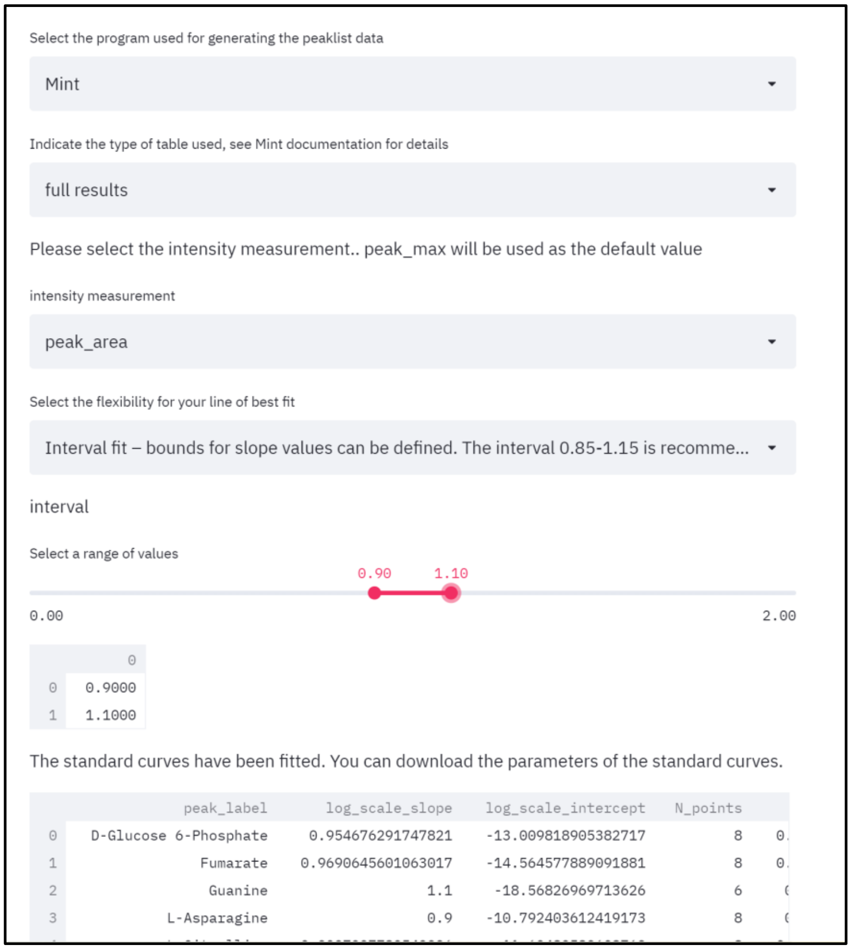
